# FGFR2 fusion protein-driven mouse models of intrahepatic cholangiocarcinoma unveil a necessary role for Erk signaling

**DOI:** 10.1101/2020.05.20.106104

**Authors:** Giulia Cristinziano, Manuela Porru, Dante Lamberti, Simonetta Buglioni, Francesca Rollo, Carla Azzurra Amoreo, Isabella Manni, Diana Giannarelli, Cristina Cristofoletti, Giandomenico Russo, Mitesh J. Borad, Gian Luca Grazi, Maria Grazia Diodoro, Silvia Giordano, Mattia Forcato, Sergio Anastasi, Carlo Leonetti, Oreste Segatto

## Abstract

**Background and aims:** About 15% of intrahepatic cholangiocarcinoma (iCCA) express fibroblast growth factor receptor 2 (FGFR2) fusion proteins (FFs), most often in concert with mutationally inactivated TP53, CDKN2A or BAP1. FFs span residues 1-768 of FGFR2 fused to sequences encoded by any of a long list (>60) of partner genes, a configuration sufficient to ignite oncogenic FF activation. In line, FGFR-specific tyrosine kinase inhibitors (F-TKI) were shown to provide clinical benefit in FF+ iCCA, although responses were partial and/or limited by resistance mechanisms, including the FF V565F gatekeeper mutation. Herein we present an FF-driven murine iCCA model and exploit its potential for pre-clinical studies on FF therapeutic targeting.

**Methods:** Four iCCA FFs carrying different fusion sequences were expressed in *Tp53*^*-/-*^ mouse liver organoids. Tumorigenic properties of genetically modified liver organoids were assessed by intrahepatic/subcutaneous transplantation in immuno-deficient mice. Cellular models derived from neoplastic lesions were exploited for pre-clinical studies.

**Results:** Tumors diagnosed as CCA were obtained upon transplantation of FF-expressing liver organoids. The penetrance of this tumorigenic phenotype was influenced by FF identity. Tumor organoids and 2D cell lines derived from CCA lesions were addicted to FF signaling via Ras-Erk, regardless of FF identity or presence of V565F mutation. Double blockade of FF-Ras-Erk pathway by concomitant pharmacological inhibition of FFs and Mek1/2 provided greater therapeutic efficacy than single agent F-TKI *in vitro* and *in vivo*.

**Conclusions:** FF-driven iCCA pathogenesis was successfully modelled in murine *Tp53*^*-/-*^ background. This model revealed biological heterogeneity among structurally different FFs. Double blockade of FF-Erk signaling deserves consideration for improving precision-based approaches against human FF+ iCCA.

Abbreviations used in this paper: ANOVA, analysis of variance; Bap1, BRCA1-Associated-Protein 1; Cdkn2a, cyclin-dependent kinase inhibitor 2A; Cftr, cystic fibrosis transmembrane conductance regulator; Ck19, cytokeratin 19; Cyp3A, cytochrome P450, family 3, subfamily A; EGF, Epidermal growth factor; EGFR, Epidermal growth factor receptor; EpCAM, epithelial cell adhesion molecule; Erk, extracellular signal–regulated kinase; FGFR2, fibroblast growth factor receptor 2; FRS2, fibroblast growth factor receptor substrate 2; GRB2, growth factor receptor-bound 2; GSEA, gene set enrichment analysis; GSVA, gene set variation analysis; H&E, hematoxylin and eosin; HepPar1, hepatocyte Paraffin 1; Hnf4α-7, hepatocyte nuclear factor 4 alpha; Hprt, hypoxanthine-guanine phosphoribosyl transferase; LGR5, leucine-rich repeat-containing G-protein coupled receptor 5; NTRK, neurotrophic Tyrosine Kinase; Parp, Poly (ADP-ribose) polymerase; RECIST, response evaluation criteria in solid tumors; SHP2, Src homology phosphatase 2; Ttr, transthyretin.

## Introduction

Intrahepatic cholangiocarcinoma (iCCA) is a neoplastic disease of biliary epithelial cells. Because of its indolent course, iCCA is often diagnosed at a locally advanced or metastatic stage. Palliative chemotherapy is the standard of care for inoperable patients, yielding a dismal 5-10% 5 year survival rate^1^.

The need for developing novel therapeutic approaches to iCCA has motivated large scale sequencing studies aimed at identifying actionable genomic drivers. Chromosomal rearrangements that give rise to FGFR2 fusion proteins were reported to occur in 12-15% of iCCA patients^2^. As depicted in Supplementary Figure 1, iCCA FGFR2 fusion proteins, henceforth referred to as FF, contain residues 1-768 of FGFR2IIIb, fused to sequences encoded by any of a long list of partner genes (>60 identified so far)^3, 4^. It is thought that fusion sequences force constitutive dimerization of adjacent FGFR2 sequences, which causes constitutive activation of the FGFR2 tyrosine kinase domain (TKD) and attendant oncogenic signaling. Whether different fusion sequences may bestow specific oncogenic properties onto FFs is a currently unresolved issue^1^.

Meaningful clinical responses were documented in FF+ iCCA patients treated with FGFR-specific tyrosine kinase inhibitors (F-TKI), including BGJ398, TAS-120 and pemigatinib, all of which have now progressed to phase 3 scrutiny as first line single agents in FF+ iCCA^1^. While providing a proof of principle that FFs are actionable iCCA oncogenic drivers, the above studies highlighted that clinical responses to FF targeting in iCCA are most often incomplete and short-lived, due to development of secondary resistance^5-7^. A number of mutations causing single aminoacid substitutions in the FF TKD were found to be a genetic determinant of secondary resistance, due to their ability to impair drug-target interactions^5, 6^. Additional routes to F-TKI resistance, likely linked also to non-genetic mechanisms, are surmised to exist in FF+ iCCA^5-7^. It therefore appears that further progress towards therapeutic targeting of FF+ iCCA requires the implementation of novel and more aggressive pharmacological approaches.

A significant hurdle to the above strategy is the lack of genetically defined iCCA models suitable for pre-clinical studies. Recent studies have shown that primary liver tumors can be modeled using liver organoids (LO)^8, 9^, i.e. 3D cultures of bipotent liver precursors^10^, which can be manipulated *in vitro* to create the desired cancer-driving genetic make-up and subsequently transplanted in mice to allow for tumor development^8, 9^. This approach was shown to yield either hepatocarcinoma (HCC) or iCCA, an outcome that reflected the specific role of different oncogenes as select drivers of either HCC (e.g. amplified *MYC*) or iCCA (e.g. mutated *KRAS*)^8^. Herein, we report on the generation and pre-clinical exploitation of a mouse model of FF-driven iCCA that capitalizes on the above described LO-based methodology.

## Materials and methods

Details on cell culture procedures and media, viral vectors, gene transfer procedures, immunoblotting, immunofluorescence, histopathology and immunohistochemistry are included in Supplementary information along with the list of reagents, antibodies, PCR primers and kits (see Supplementary Tables 3-6).

### Animal studies

All animal experimentation procedures were approved by the ethics committee of the Regina Elena National Cancer Institute (code 12048) and the Italian Ministry of Health (code 947/2015-PR) and were in compliance with the national and international directives (D.L. March 4, 2014, no. 26; directive 2010/63/EU of the European Parliament and European Council; Guide for the Care and Use of Laboratory Animals, United States National Research Council, 2011).

Male NOD-SCID mice (4-6 weeks old) were purchased from Charles River Laboratory. For tumorigenicity assays, organoids were infected with a luciferase-expressing lentivirus (LV) before being transplanted (5×10^5^ cells/animal) either in the liver or subcutaneous (s.c.) tissue of recipient mice. Intrahepatic tumor growth was assessed by live luciferase detection using IVIS Lumina II CCD camera system (Perkin Elmer). Data were acquired and analysed using the Living Image Software version 4.3 (Perkin Elmer). Photon emission was measured in specific regions of interest (ROIs). Data were expressed as photon/second/cm^2^/steradian (p/s/cm^2^/sr). The intensity of bioluminescence was color-coded for imaging purposes. For s.c. tumorigenicity assays tumor growth was followed by caliper measurements three times per week, using the following formula: volume = (width^2^ x length)/2. When required, mice were anesthetized and euthanized by cervical dislocation.

Secondary s.c. tumors for pharmacological studies were obtained by transplantation of F-BICC1 tumoroids (Tum 1, 2×10^5^cells per s.c. site) in NOD-SCID mice. Tumors reached a volume of 100-130 mm^3^ 12-14 days past tumoroids implantation, at which time mice were randomized to receive drug treatment. In experiment #1 (presented in Figure 6), BGJ398 and trametinib were dosed by oral gavage at 15 mg/kg and 1 mg/kg respectively in single agent and B+T groups. In experiment #2 (presented in Figure 7), single agent BGJ398 and trametinib were dosed as in experiment #1. In the B+T group the dose of both BGJ398 and trametinib was reduced by 20% (i.e. 12 mg/kg BGJ398 and 0.8 mg/kg trametinib). On day 15, we subdivided the B+T mice into two different subgroups, based on tumor volume and body weight loss: B+T-1 group (n = 5) received no further drug treatment, while B+T-2 group (n = 5) was treated for two additional days in the third week (i.e. day 16 and day 19). Likewise, on day 15, mice in the BGJ398 group were randomized to two subgroups: B/B+T-1 received B+T twice a week (i.e. day 16 and day 19); B/B+T-2 received B+T for 5 days/week.

Tumor growth was monitored by caliper measurement. Tumor volume values were also used for assessing treatment efficacy via response evaluation criteria in solid tumors (RECIST 1.1) criteria adapted to animal experimentation^11^. Thus, any tumor volume change ≥35% from baseline was rated as progressive disease (PD), any tumor volume change comprised between -50% and +35% from baseline was rated as stable disease (SD) and any tumor volume reduction ≥50% from baseline was rated as partial response (PR).

### Statistical analysis

Values were expressed as average ± SEM. One-way analysis of variance (ANOVA) with a post hoc Bonferroni’s test was used for multiple sample comparisons. The student’s t-test (unpaired, two-tailed) was used for single pair-wise comparisons. Differences were considered statistically significant when P <0.05. Data were elaborated using GraphPad Prism Software 8.0.

## Results

### *Tp53* null mouse liver organoids transduced with FGFR2-BICC1 or FGFR2-TACC3 give rise to iCCA when transplanted in immuno-compromised mice

In iCCA, *FGFR2* fusions have been shown to co-occur with mutations affecting *TP53, CDKN2A* or *BAP1*^12, 13^, which likely reflects a requirement for loss of defined tumor suppressor pathway/s during FF-driven iCCA pathogenesis. Of note, *TP53* mutations were reported to be a negative prognostic factor for FF+ iCCA patients^12^. With this in mind, we chose to model FF-driven tumorigenesis in a *Tp53* null background and therefore used livers explanted from *Tp53*^*-/-*^ C57BL/6J mice (Supplementary Figure 2A) to generate organoids. In line with previous reports, we observed that organoids developed as buds extending from biliary ducts^10, 14^. Upon passaging, these outgrowths yielded homogeneous cultures of organoids having the aspect of spherical cysts, as described^10, 14^ (Figure 1A).

**Figure 1.**
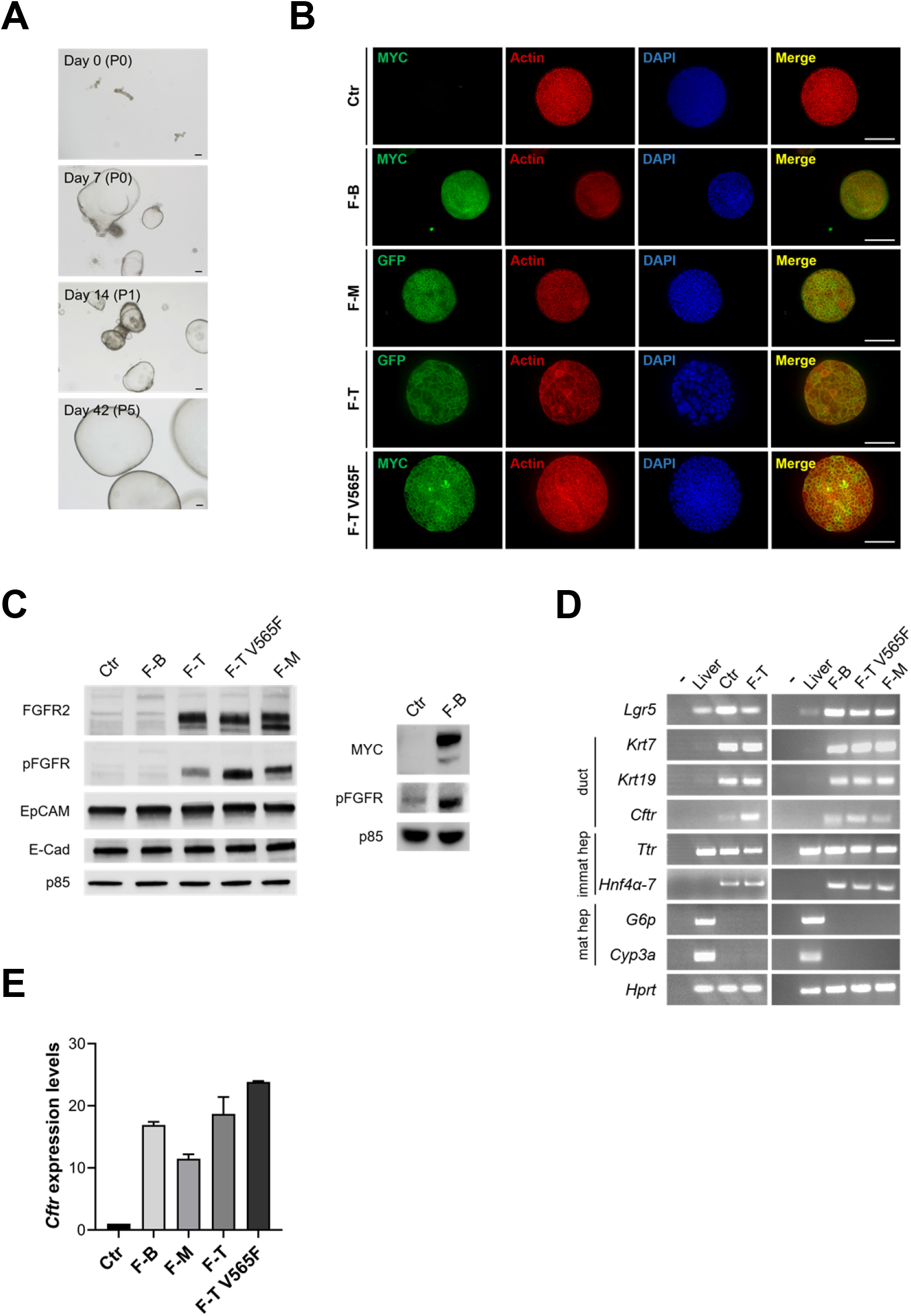
Generation of liver organoids from *Tp53*^-/-^ mice. (*A*) Ductal structures obtained after enzymatic digestion of *Tp53*^-/-^ mice livers were seeded in matrigel (day 0). Ductal outgrowths were visualized after a few days and generated organoids at later passages (P). Scale bar: 100 μm. (*B*) Organoids transduced with control (Ctr) and FF-encoding recombinant retrovirus were imaged by anti-MYC tag immunostaining or GFP fluorescence (green); actin was visualized by phalloidin (red) and nuclei by DAPI stain. Scale bar: 100 μm. (*C*) Cell lysates obtained from the indicated organoids (top) were probed with the indicated antibodies. Because steady state expression levels of F-BICC1 were lower than those of other FFs, unambiguous detection of F-BICC1 required loading of more lysate and blotting with anti-MYC antibody (right panel). (*D*) Transcripts encoding the indicated markers were assessed by RT-PCR analysis of liver organoids infected with Ctr or FF-expressing retrovirus stocks. RNA obtained from *Tp53*^-/-^ liver was used as control. Duct: biliary duct marker; immat hep: immature hepatocyte marker; mat hep: mature hepatocyte marker. (E) Expression levels of the mature bile duct marker *Cftr* in the indicated organoids were assessed by quantitative real time PCR analysis.

Three structurally different FFs previously identified in iCCA, namely FGFR2-BICC1, FGFR2-MGEA5 and FGFR2-TACC3 (henceforth abbreviated as F-BICC1, F-MGEA5 and F-TACC3 in the main text or F-B, F-M and F-T in figures)^15^ were expressed in *Tp53*^*-/-*^ liver organoids by retrovirus-mediated gene transfer. We also generated virus stocks for expressing the BGJ398-resistant F-TACC3 V565F mutant^16^. Empty retrovirus was used as control. Puromycin-selected organoids consisted of cells positive for either GFP fluorescence (GFP-tagged F-MGEA5 and F-TACC3) or anti-MYC 9E10 immunoreactivity (MYC-tagged F-BICC1 and F-TACC3 V565F) (Figure 1B). Both imaging methods outlined the cell periphery (see Supplementary Figure 2B for higher resolution), in agreement with FFs being exposed at the cell surface^16^. Organoids infected with control virus scored negative. Immunoblot analysis confirmed expression and constitutive phosphorylation of FFs in transduced organoids, with F-BICC1 being expressed at lowest levels (Figure 1C). In line with their stem-like bipotent differentiation potential, control organoids expressed the stem cell marker *Lgr5* along with markers of both hepatocyte and cholangiocyte lineages^17^. This molecular profile was not appreciably altered by FF expression (Figure 1D). Interestingly, we detected increased expression of the mature cholangiocyte marker *Cftr*^18, 19^ in FF-transduced organoids (Figure 1D), a datum that was confirmed by quantitative real time PCR (Figure 1E).

In order to test their tumorigenic potential, control and FF-expressing organoids were transplanted in NOD-SCID mice, either in the liver sub-capsular region or subcutaneously (s.c.)^8^. Growth of orthotopic and s.c. allotransplants was monitored by longitudinal luciferase bioimaging and caliper measurement, respectively. Control organoids were not tumorigenic, in agreement with previous reports^8, 9^. Instead, allotransplants of F-BICC1 and F-TACC3 organoids yielded progressively growing tumors at either orthotopic or s.c. transplantation sites (Figure 2A, B), with 100% penetrance (Supplementary Table 1). Growth of liver tumors led to progressive health deterioration of recipient mice, eventually requiring their euthanasy. At necropsy, liver tumors appeared as nodular masses deforming the liver surface (Figure 2C and Supplementary Figure 3A). Liver tumors were diagnosed as moderately to poorly differentiated adenocarcinomas (Figure 2D and Supplementary Figure 3B) containing areas of dense stromal reaction, visualized by Masson’s trichromic stain (Figure 2D and Supplementary Figure 3B), as typically observed in iCCA. Tumor lesions stained positive for the bile duct marker Ck19, while being negative for the hepatocyte marker HepPar1 and contained cycling Ki67+ neoplastic cells. Expression of FGFR2 fusions was confirmed by either anti-MYC tag or anti-FGFR2 stain (Figure 2D and Supplementary Figure 3B). In MYC+ tumors, stromal cells were negative for anti-MYC reactivity, which implied their host origin (Figure 2D). Tumors that developed at s.c. injection sites appeared as light grey nodules with a hard texture (Figure 2C and Supplementary Figure 3A) and were histologically and immuno-phenotypically similar to the above described liver neoplastic lesions (Figure 2E and Supplementary Figure 3C).

**Figure 2.**
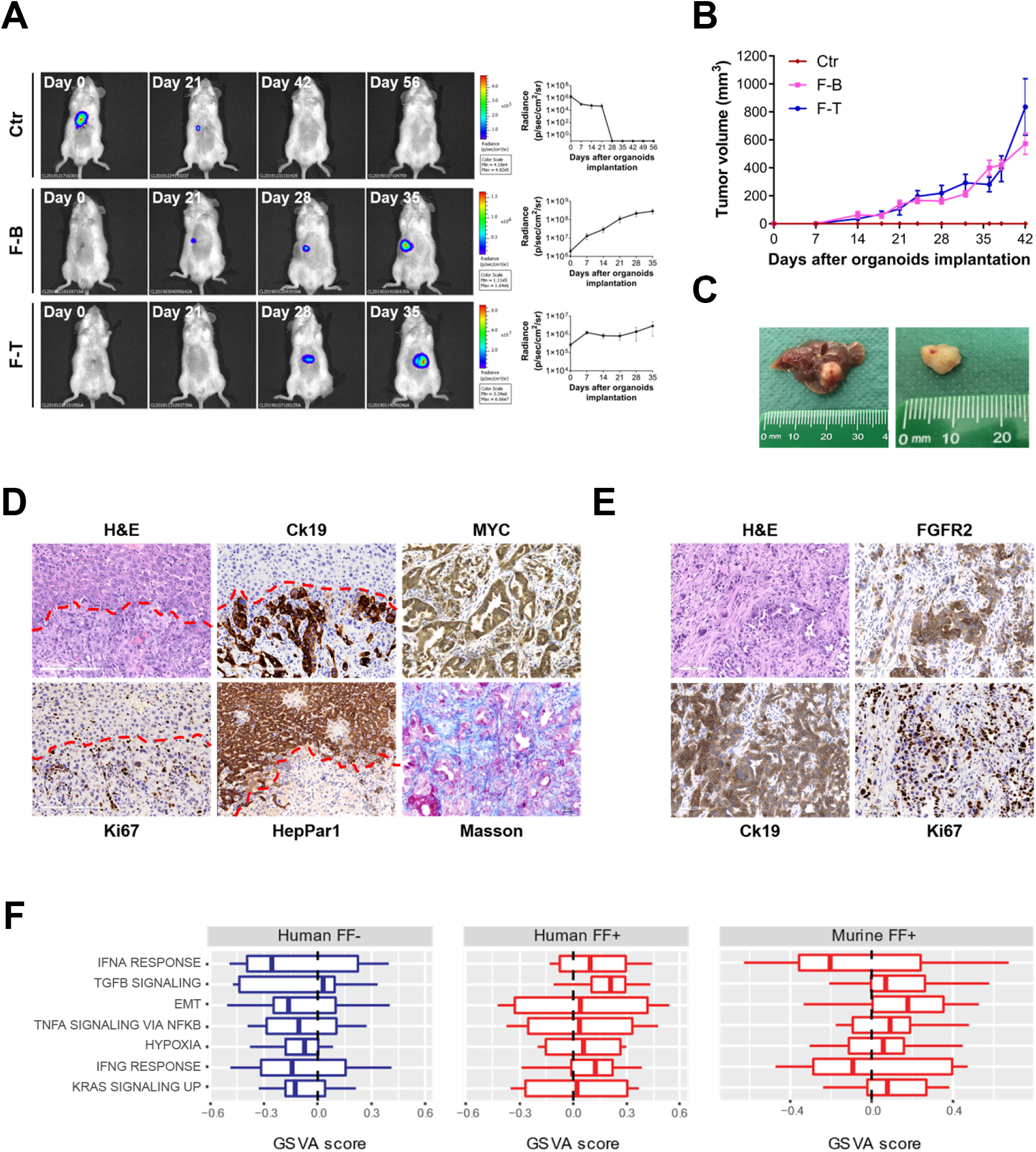
Tumorigenicity of liver organoids expressing F-BICC1 or F-TACC3. (*A*) Growth of liver implants of Ctr, F-BICC1 and F-TACC3 organoids was imaged by longitudinal live measurements of luciferase activity. The intensity of bioluminescence was color-coded for imaging purposes and plotted over time. (*B*) Growth of tumors derived from s.c. implantation of Ctr and FF-expressing organoids was monitored by longitudinal caliper measurement. (*C*) Representative images of liver (left) and subcutaneous (right) tumors obtained upon transplantation of F-BICC1 liver organoids. (*D*) Histopathology of liver tumors obtained upon intrahepatic transplantation of F-BICC1 organoids. Tissue sections were stained as indicated. Red dotted lines demarcate the boundary between tumor and normal liver. The MYC stain identifies MYC-tagged F-BICC1. Note that collagen fibers appear as blue-colored areas in Masson’s trichrome stain. (*E*) Histopathology of tumors obtained upon s.c. transplantation of F-BICC1 liver organoids. Sections were stained as indicated. (*F*) Gene set enrichment of RNA-seq profiles from human iCCA (left and middle plots) and murine iCCA driven by either F-BICC1 or F-TACC3 (right plot). Boxplots depict GSVA activity scores of significantly upregulated gene sets in FF+ versus FF-human iCCA. Red boxes refer to FF+ tumors (whether human or murine), blue boxes indicate FF-human iCCA.

F-MGEA5 expressing organoids were not tumorigenic when transplanted in the liver, even after a 190-day follow-up (Supplementary Figure 4A) and yielded *bona fide* iCCA-like lesions in only 1/8 s.c. injection sites (Supplementary Figure 4B and Supplementary Table 1). This tumor had histopathological features similar to those observed in F-BICC1 and F-TACC3 lesions (Supplementary Figure 4C). In the remaining seven s.c. transplantation sites we observed the development of cystadenomas lined by dysplastic epithelium. These cystic lesions lacked signs of overt neoplastic transformation, except in one case in which we observed an area showing clear progression to cystic adenocarcinoma (Supplementary Figure 4D). We note that the low tumorigenic potential of F-MGEA5-expressing organoids was unlikely to be caused by poor expression/activation of F-MGEA5 itself, which, in fact, was as good as that of F-TACC3 (Figure 1C). Furthermore, the expression/activation of F-MGEA5 in the single iCCA-like tumor we obtained was comparable to that of F-TACC3-driven tumors (Supplementary Figure 4E, see also Figure 3A), implying that F-MGEA5-dependent tumorigenicity did not require selection for super-high level F-MGEA5 expression/activation.

**Figure 3.**
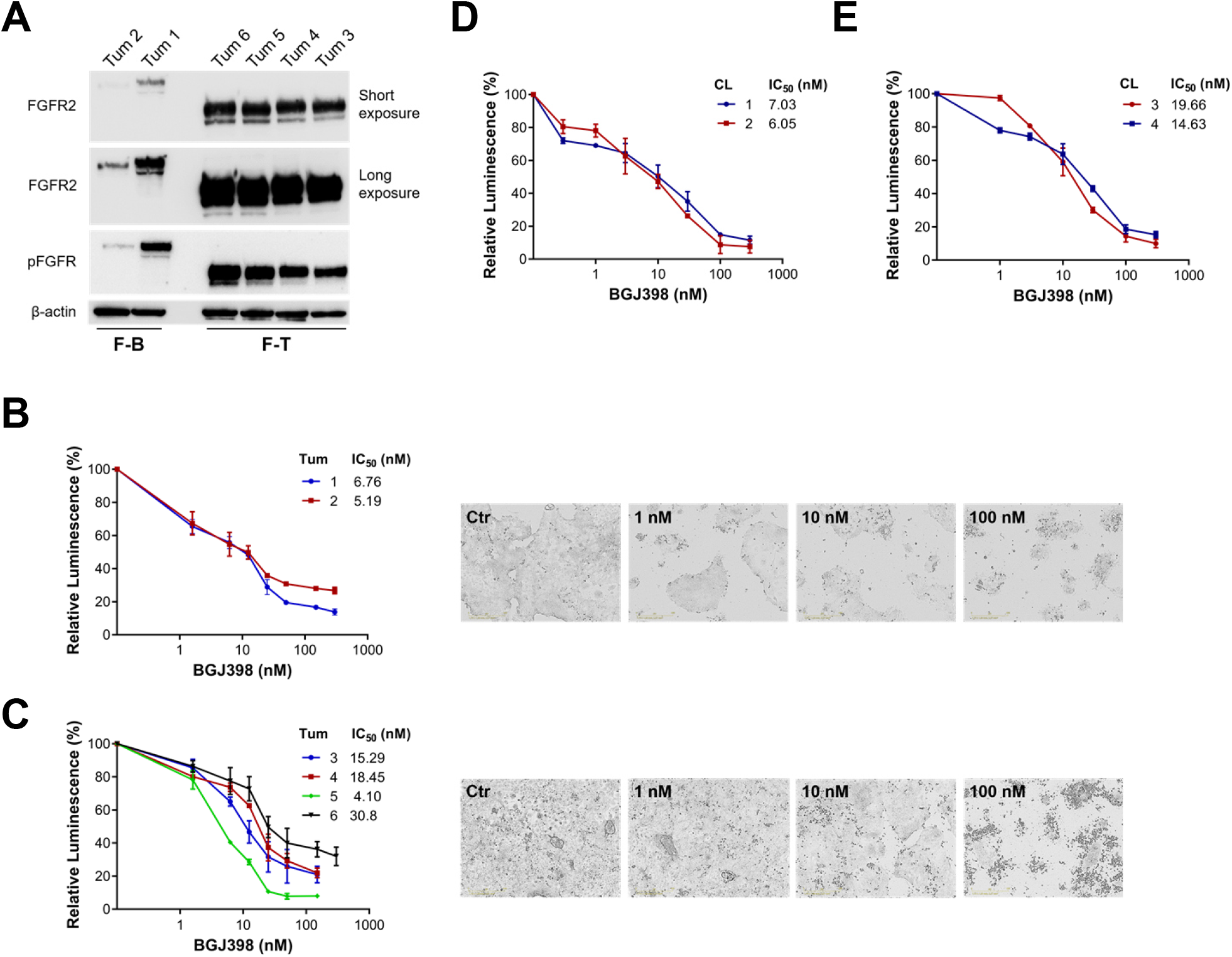
Cellular models derived from iCCA lesions are addicted to FF signaling. (*A*) Analysis of expression and activation of F-BICC1 and F-TACC3 in tumoroids derived from iCCA like lesions. Tum 2 and Tum 3-5 were derived from liver lesions, Tum 1 and 6 from s.c. lesions. Total cell lysates were probed with the indicated antibodies. (*B, C*) Dose-dependent profile of growth inhibition in BGJ398-treated F-BICC1 (B) and F-TACC3 (C) tumoroid cultures. Cell growth was assayed after 72 hours of drug treatment using the CellTiter-Glo kit and plotted as % decrease of luminescence over untreated cells. IC_50_ values are indicated for each tumoroid. Representative photographs of Tum 1 (B) and Tum 4 (C) were taken at the 72-hour endpoint using Incucyte S3 Live-Cell Imaging System. (*D, E*) Dose-dependent profiles of growth inhibition in BGJ398-treated 2D cell lines derived from F-BICC1 (D: CL 1 and CL 2) and F-TACC3 (E: CL 3 and CL 4).

Our gene set enrichment analysis (GSEA) of RNA-seq data from the iCCA TCGA cohort^20^ revealed that a number of transcriptional signatures were significantly up/down-regulated in FF+ compared to FF-samples (Supplementary Table 2). Notably, FF+ iCCA samples were associated with higher activity of well-known cancer-relevant pathways, including KRAS signaling and epithelial-mesenchymal transition. By single sample gene set variation analysis (GSVA) we measured the activity of these up-regulated pathways in RNA-seq profiles obtained from murine iCCA lesions and, reassuringly, observed a similar picture, except that our murine iCCA models did not show prominent up-regulation of interferon-dependent signatures (Figure 2F). Most likely, this is imputable to murine tumors having thrived in the NOD-SCID immuno-deficient context^21^. In passing, we note that interferon signaling is a prominent driver of cancer immuno-editing^22^, which raises the question whether immunotherapeutic approaches could be effective against FF+ iCCA. We also interrogated our RNA-seq data to search for possible differences between iCCA lesions driven by different FFs. Interestingly, F-BICC1- and F-TACC3-driven murine tumors tended to group separately from each other (Supplementary Figure 3D). In aggregate, these data point to a) relevant similarities between our murine model and human FF+ iCCA as a whole; b) a degree of inter-FF differences, potentially causative of biological diversification.

### Cellular models derived from iCCA lesions are addicted to oncogenic signaling by FGFR2 fusions

We derived tumor organoids (henceforth referred to as tumoroids)^23^ and conventional 2D cell lines from liver and s.c. tumors expressing F-BICC1 and F-TACC3. Each tumoroid expressed the expected FF, which displayed constitutive catalytic activation, as assessed by WB analysis with an antibody recognizing the pY657/pY658 motif required for full FGFR2 catalytic activation^24^ (Figure 3A). Cell lines growing in 2D expressed the expected FGFR2 fusions along with the Ck19 biliary cell marker (Supplementary Figure 5A). As already observed in transduced organoids, F-BICC1 expression was lower than that of F-TACC3 also in tumoroids and 2D cell lines. Notably, F-BICC1 displayed a higher stoichiometry of Y657/Y658 phosphorylation compared to F-TACC3 (Figure 3A and Supplementary Figure 5B), suggesting that its relatively low expression was compensated for by a high level of intrinsic tyrosine kinase activity.

The clinically advanced BGJ398 F-TKI^25^ caused dose-dependent suppression of tumoroid growth/viability (Figure 3B, C), with IC_50_ values in the low nM range. Similar results were obtained with 2D cell lines (Figure 3D, E and Supplementary Figure 5C). We conclude that the above 2D and 3D cellular models are dependent on FF signaling, possibly reflecting the state of oncogenic addiction of their tumor of origin.

Next, we investigated signaling cascades downstream to FFs in iCCA cell lines. We focused on the Ras-Erk pathway, because previous work highlighted its potential relevance downstream to FFs^6^ and our RNA-seq data pointed to upregulation of the Kras-driven transcriptional program in human and mouse FF+ iCCA. The docking/scaffolding Frs2 adaptor protein was phosphorylated on tyrosine residues responsible for the recruitment of Grb2 and Shp2^26^, as indicated by anti-pFrs2 reactivity in immunoblot analyses of cell lysates (Figure 4A and Supplementary Figure 6A) and pull-down assays employing GST-GRB2 SH2 and GST-SHP2 (N+C) SH2 baits (Supplementary Figure 6B). Shp2 was phosphorylated on Tyr residues involved in Grb2 binding, as indicated by anti-pSHP2 reactivity in cell lysates (Figure 4A and Supplementary Figure 6A) and GST-GRB2 SH2 pull-down assays (Supplementary Figure 6C). Tyr phosphorylation of Frs2 and Shp2 correlated with the activating phosphorylation of Mek1/2 and Erk1/2 (Figure 4A and Supplementary Figure 6A). All of the above phosphorylation events required FF catalytic activity, because they were suppressed by BGJ398 (Figure 4A and Supplementary Figure 6A-C). These data fit in the currently accepted model of FGFR2 activation^27^ and support that FFs drive Ras-Erk activation by instructing formation of activity-dependent molecular complexes involving Frs2, Shp2 and Grb2^27^. In line with this model, pharmacological inhibition of Shp2 by SHP099^28^ caused a remarkable reduction of Erk1/2 activation downstream to both F-BICC1 and F-TACC3 (Figure 4B). It is notable that inhibition of the FF-Erk1/2 signaling axis by BGJ398 was durable, with no apparent evidence of late pathway reactivation (Figure 4A and Supplementary Figure 6A). The protracted signal inhibition achieved by BGJ398 in the above cellular models is congruent with the potent activity exhibited by BGJ398 in cell viability/growth assays (Figure 3B-E).

**Figure 4.**
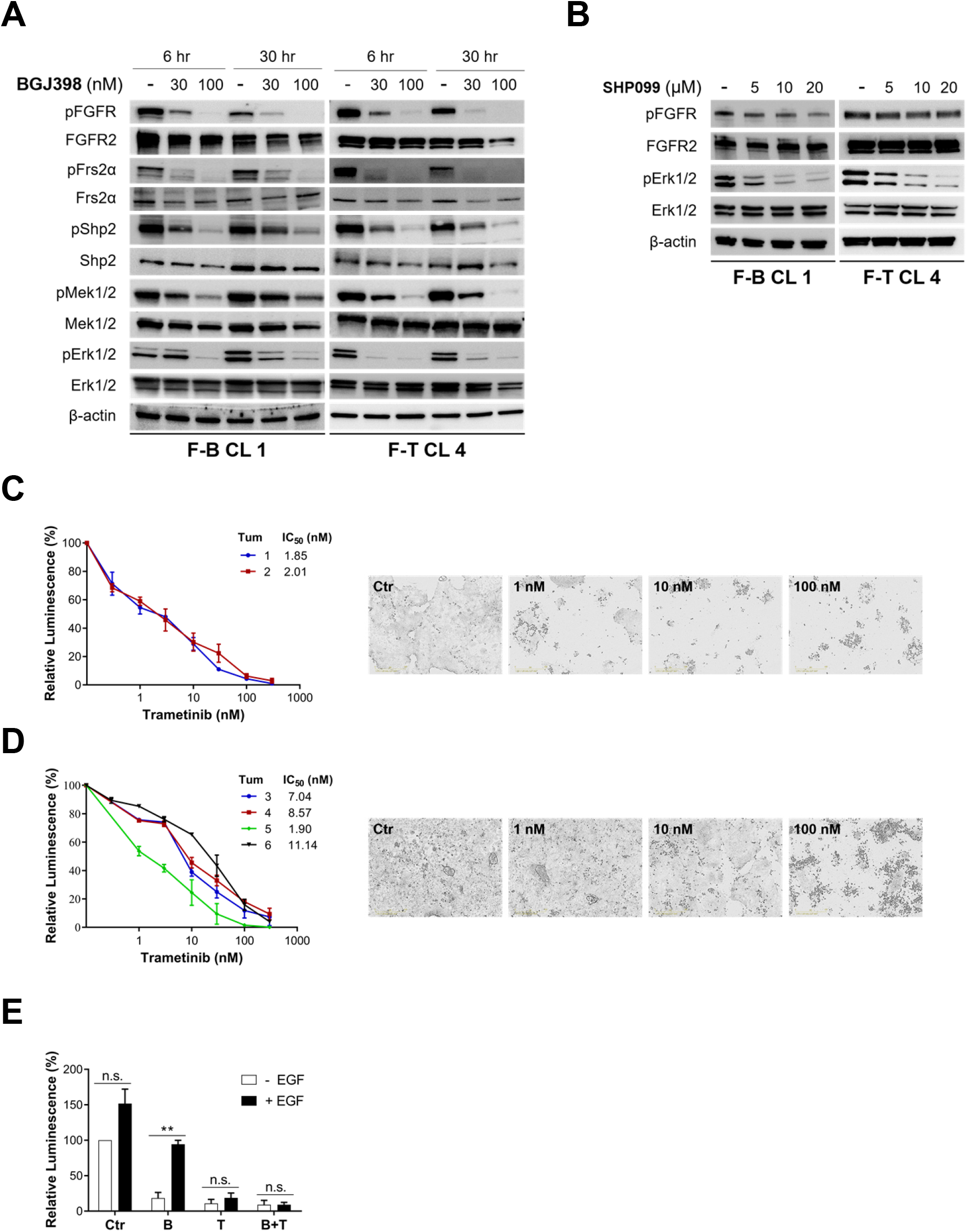
FF-driven iCCA cells are dependent on downstream Ras-Erk activity. (*A*) F-BICC1 and F-TACC3 iCCA cell lines were treated with the indicated concentrations of BGJ398 for 6 and 30 hours before lysis. Total cell extracts were immunoblotted as indicated. (*B*) F-BICC1 and F-TACC3 iCCA cell lines were incubated for 4 hours with the SHP2 inhibitor SHP099. Total cell extracts were immunoblotted as indicated. (*C, D*) F-BICC1 (C) and F-TACC3 (D) tumoroids were exposed to escalating concentrations of trametinib for 72 hours. Plots indicate growth suppression (expressed as % decrease of luminescence over untreated control cells) as determined by the CellTiter-Glo kit. IC_50_ values are indicated for each tumoroid. Representative photographs of Tum 1 (C) and Tum 4 (D) were taken at the 72-hour time-point of growth assays using Incucyte S3 Live-Cell Imaging System. (E) F-TACC3 tumoroid cells (Tum 3) were grown -/+ EGF (10 ng/ml) in presence or absence of BGJ398 (B, 25 nM), trametinib (T, 25 nM) or their combination (B+T). Cell growth was measured with the CellTiter-Glo kit. Note that EGF enhanced cell proliferation in control untreated cells. See also Supplementary Figure 7E. Data are presented as mean ± SEM. One-way analysis of variance (ANOVA) with a post hoc Bonferroni’s test was used to evaluate statistical significance among the indicated treatments (\*\**P* = 0.009).

### Erk1/2 is a necessary signaling hub downstream to FGFR2 fusions in iCCA

Trametinib^29^, a clinically approved MEK1/2 inhibitor, suppressed growth/viability of iCCA tumoroids (Figure 4C, D) and 2D cell lines (Supplementary Figure 7A, B), in fact phenocopying BGJ398. In line, the SHP2 inhibitor SHP099 was also effective in curbing growth/viability of F-BICC1-addicted iCCA cells (Supplementary Figure 7C, D). We also generated an iCCA model driven by the BGJ398-resistant FGFR2-TACC3 V565F^5^ mutant (Supplementary Figure 8A-D and Supplementary Table 1) and determined that F-TKI resistant tumoroids (Supplementary Figure 8E, F) still required downstream Erk1/2 signaling (Supplementary Figure 8G).

The above data support a model whereby Erk1/2 activation plays a necessary role in shaping addiction of iCCA cells to FF oncogenic signaling. If so, reactivation of signal flow through the Ras-Erk pathway should rescue FF-driven iCCA cells from BGJ398 inhibition. Indeed, epidermal growth factor (EGF) was able to alleviate BGJ398-mediated growth inhibition in F-TACC3 tumoroids. Addition of trametinib nullified EGF-dependent rescue (Figure 4E and in Supplementary Figure 7E), implying that the EGF signal rescued BGJ398-treated cells through bypass activation of Ras-Erk.

### Oncogenic signaling of FGFR2 fusions in iCCA cells is highly sensitive to combined blockade of FGFR2 fusions and Mek1/2

Concomitant targeting of multiple components of a signaling pathway engaged by an oncogenic driver has been shown to be more efficacious than blockade of the oncodriver in isolation^30, 31^. Therefore, we hypothesized that the efficacy of FF targeting could be improved by combining FF blockade with Mek1/2 inhibition. As shown in Figure 5A, B, BGJ398 and trametinib interacted synergistically in short-term growth assays of F-BICC1 and F-TACC3 tumoroids. This was confirmed in long term growth assays of F-BICC1 and T-TACC3 cell lines (Figure 5C). At the biochemical level, synergy between BGJ398 and trametinib correlated with stronger inhibition of cyclin D1 expression and marked induction of cleaved Parp1 (Figure 5D).

**Figure 5.**
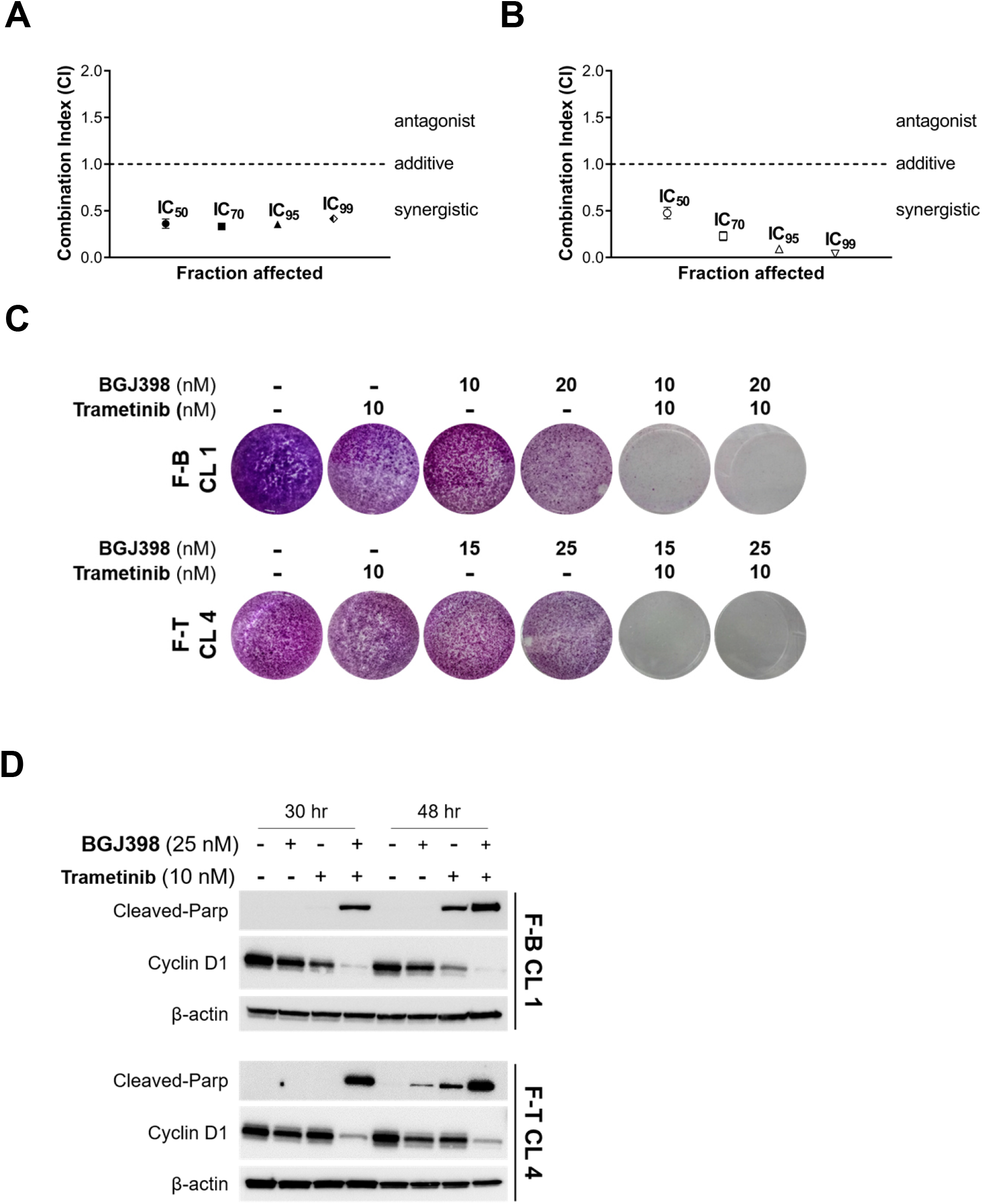
Improved targeting of FF oncogenic dependence by combined FF and Mek1/2 blockade. (*A, B*) The combination index (CI) for the BGJ398 + trametinib combination was calculated according to Chou-Talalay, as detailed in Materials and Methods. Values <1 indicate synergistic drug interaction. A and B show data obtained with Tum 1 (F-BICC1) and Tum 4 (F-TACC3), respectively. (*C*) F-BICC1 (top) and F-TACC3 (bottom) cell lines were treated with BGJ398 and trametinib, either as single agents or in combination. Cells were stained with crystal violet. (*D*) Semi-confluent monolayers of F-BICC1 and F-TACC3 cell lines were treated with the indicated drugs for 30 or 48 hours before lysis. Total cell lysates were immunoblotted as indicated.

### Therapeutic targeting of FGFR2-BICC1-driven iCCA in the mouse by the BGJ398 + trametinib combination

To validate *in vivo* the above results, we set out to compare the efficacy of single agent BGJ398 or trametinib versus the BGJ398 + trametinib (B+T) combination in immuno-deficient mice carrying tumors generated by subcutaneous injection of F-BICC1 tumoroids. In mice treated with vehicle, F-BICC1 tumors continued to grow rapidly, reaching an average volume of approximately 700 mm^3^ after 12 days (Figure 6A), at which time mice were euthanized because of severe skin ulceration at the site of tumor growth. Single agent trametinib provided only for short-lived delay of tumor growth (which was not unexpected given that MEK1/2 inhibitors reportedly have limited therapeutic efficacy as single agents^29^), with euthanasy being necessary by day 15. In mice treated with BGJ398, tumors underwent sizeable shrinkage until day 12, but resumed growth thereafter and progression continued unabated until day 22 (Figure 6A). Remarkably, the B+T combination showed superiority to single agent BGJ398 already at the day 15 time-point and remained significantly better than single agent BGJ398 until day 22 (Figure 6A). This was consistent with pharmacodynamic studies showing that the B+T combination elicited stronger inhibition of pErk immunoreactivity when compared to single agents (Figure 6C; note that lack of suitable reagents prevented evaluation of pFGFR, see Supplementary Figure 9). When RECIST 1.1-like criteria adapted to experimentation in the mouse^11^ (see Methods for details) were used for evaluating responses of individual mice, we noted that a partial response (PR) was reached at nadir in 8/8 (100%) tumors in the B+T group, which contrasted with PR being observed as best response in only 3/7 tumors (42%) in the BGJ398 group (Figure 6B). However, two sudden deaths occurred in the B+T group between day 20 and 21, despite the fact that daily animal inspection and regular body weight measurements (Supplementary Figure 10A) had not raised suspicion of overt toxicity. Previous studies had reported similar dosing of the B+T combination in mice, with no evidence of lethal events^32, 33^.

**Figure 6.**
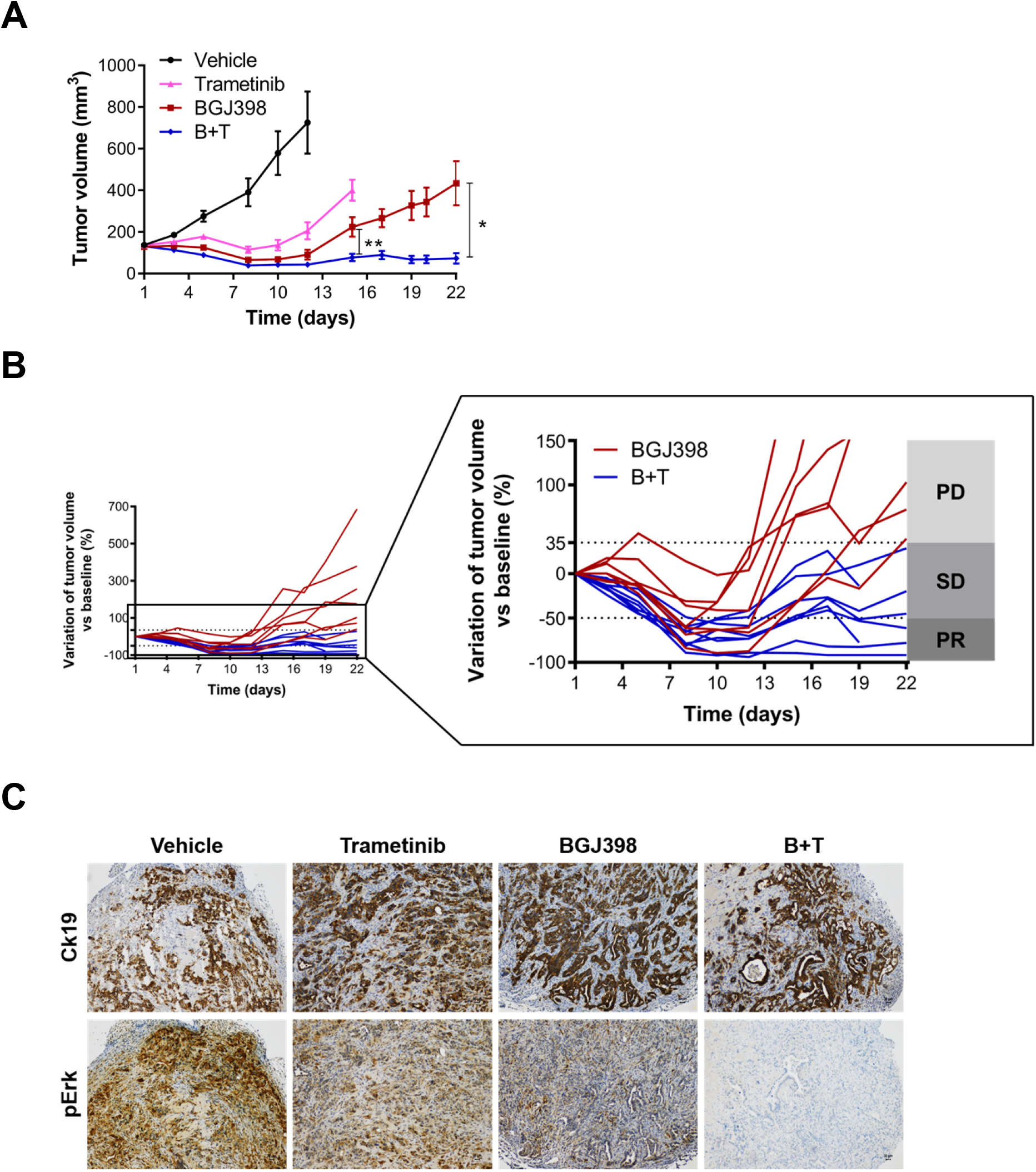
Therapeutic targeting of F-BICC1 tumors *in vivo*. iCCA-like lesions were generated by s.c. transplantation of F-BICC1 tumoroid 1. (*A*) Trametinib (1 mg/k) and BGJ398 (15 mg/kg) were administered for 5 days per week either as single agents or in combination. Control mice were treated with vehicle. Average tumor volumes from each experimental group were plotted over time. Groups were composed as follows: vehicle n = 7, trametinib n = 7, BGJ398 n = 7, B+T n = 8. Data are presented as mean ± SEM. Two-tailed unpaired t-test was used to evaluate significant differences between BGJ398 and B+T groups: P = 0.0089 on day 15, P = 0.0106 on day 22. (*B*) The spaghetti plot shows individual responses (tumor volume changes) in mice assigned to either BGJ398 (dark red) or B+T (dark blue) treatment. The enlarged plot allows for evaluation of individual responses according to RECIST 1.1-like criteria at higher resolution. PD, progressive disease (≥35% increase from baseline); PR, partial response (≤50% reduction from baseline); SD: stable disease (intermediate changes from baseline). Note that two deaths occurred on day 20-21 in the B+T group; these mice were evaluated until day 19. (*C*) Pharmacodynamic evaluation of drug efficacy in tumors treated with placebo or the indicated drugs every 24 hours for two days and explanted three hours after the last drug treatment. Tumor cells were stained with anti-Ck19 stain (top). Anti-pErk immunoreactivity was used as surrogate marker to assess drug activity. Scale bar: 30 μm.

Because adverse events in the B+T group imposed a degree of caution in the interpretation of the above experiment (henceforth referred to as experiment #2, to distinguish it from experiment #1 presented in Figure 6), we carried out a second experiment in which the dose of both BGJ398 and trametinib was reduced by 20% in the B+T group; furthermore, B+T was administered according to a de-escalation protocol (5 days/week for the first week, 4 days/week for the second week). In line with experiment #1, we observed rapid tumor growth in the control group and marginal benefit from single agent trametinib (Figure 7A). Again, the B+T combination outperformed single agent BGJ398 at the day 15 time-point (Figure 7A), which translated into 100% of B+T tumors achieving a PR at nadir, compared to 60% in the BGJ398 group (Figure 7B). Interestingly, histopathological evaluation of tumors explanted on day 15 showed that tumor responses to BGJ398 and B+T treatment were associated with the acquisition of a more differentiated glandular phenotype and increased fibrosis, particularly in B+T tumors (Figure 7C), when compared to carrier- or trametinib-treated tumors.

**Figure 7.**
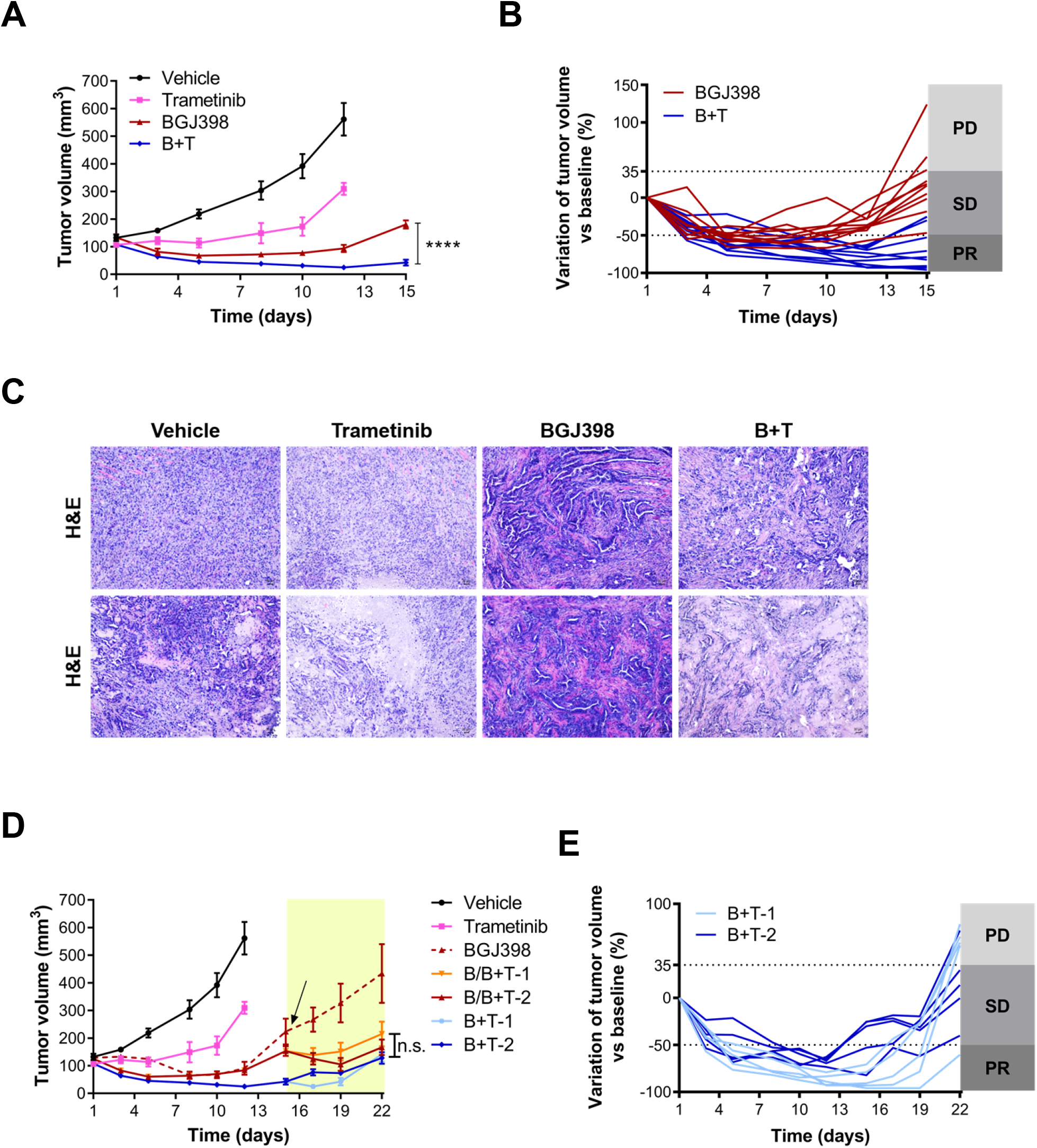
Therapeutic targeting of F-BICC1 tumors *in vivo*. (*A*) Trametinib (1 mg/kg) and BGJ398 (15 mg/kg) were administered to tumor bearing mice for 5 days per week as single agents. For the B+T combination, BGJ398 was dosed at 12 mg/kg and trametinib was dosed at 0.8 mg/kg, using the de-escalation protocol described in the text. Average tumor volumes from each experimental group were plotted over time. Groups were composed as follows: vehicle n = 6, trametinib n = 6, BGJ398 n = 10, B+T n = 10. Data are presented as mean ± SEM. Two-tailed unpaired t-test was used to evaluate significant differences between BGJ398 and B+T groups (P <0.0001 on day 15). (*B*) The spaghetti plot shows individual responses (tumor volume changes), evaluated according to RECIST 1.1-like criteria, in mice assigned to either BGJ398 (dark red) or B+T (dark blue) treatment. (*C*) Tumors explanted on day 15 from mice that underwent to the indicated drug treatments were stained with H&E. Top and bottom rows contain images from two independent tumors. Scale bar: 30 μm. (*D*) Adaptive therapeutic treatment of mice belonging to the B and B+T groups (same as in 7A), which were assigned to the indicated subgroups during the third week of the experiment (highlighted in the yellow area). Vehicle and trametinib groups are the same as in panel 7A and are reported here for reference. The dark red dotted line refers to single agent BGJ398 group in experiment #1, i.e. it corresponds to the dark red solid line in Figure 6A. Groups: B/B+T-1 n = 5, B/B+T-2 n = 5, B+T-1 n = 5, B+T-2 n = 5. Data are presented as mean ± SEM. One-way analysis of variance (ANOVA) with a post hoc Bonferroni’s test was used to evaluate significant differences in the tumor volumes between the indicated subgroups. Differences among the four subgroups in the yellow-shaded area lacked statistical significance (P >0.05 on day 22). (*E*) RECIST 1.1-like evaluation of individual tumor responses in mice assigned to the B+T-1 (light blue) and B+T-2 (dark blue) subgroups.

Aiming to prolong therapeutic benefit beyond the day 15 time-point, we opted for an adaptive trial strategy for reasons detailed below. Average tumor growth on day 15 in the BGJ398 group was comparable to that observed in experiment #1 and therefore indicative of therapeutic resistance (Figure 7D: the dark red dotted line corresponds to the BGJ398 group from Figure 6A and is reported as reference). Given the superior activity of the B+T combo, these mice were randomized to receive B+T (Supplementary Figure 10B), either twice a week (B/B+T-1) or five days a week (B/B+T-2). B+T provided therapeutic benefit in both subgroups, so that tumor growth levelled off and at day 22 ended up approaching that of mice receiving B+T *ab initio* (Figure 7D: compare the two B/B+T subgroups with the BGJ398 group in experiment #1, which is indicated by the dark red dotted line; see also Supplementary Figure 10D, E).

Looking at mice in the B+T group, we were concerned by the observation that on day 12, i.e. at the end of the second B+T cycle, five mice had experienced an average weight loss value of 8.5% (henceforth referred to as B+T-1 subgroup, Supplementary Figure 10C). Interestingly, in each of these mice the tumor volume had regressed by more than 70% (range 70.6-95.9%) (Supplementary Figure 10B). The remaining five mice (B+T-2 subgroup) experienced more contained weight loss (2.5% average value, Supplementary Figure 10C). In this subgroup, tumor shrinkage ranged between -82.3% and -25.2%, with only a single animal showing a tumor volume reduction >70% (Supplementary Figure 10B). Considering their overall performance status, mice in the B+T-1 subgroup received no further therapy. The B+T-2 subgroup was assigned to continue B+T twice a week. Tumor growth remained under control in both groups until day 19 (which in the B+T-1 subgroup corresponded to a remarkable eight-day drug-free interval), but it resumed thereafter (Figure 7D, E). The experiment was stopped on day 22, because most mice had developed skin ulcerations at the tumor site. Looking at the overall performance of the four subgroups exposed to different schedules of B+T treatment in comparison to single agent BGJ398, it is clear that the B+T combo afforded better and more prolonged control of tumor growth also in experiment #2, in which no deaths were recorded.

## Discussion

Herein, we present the generation of an iCCA mouse model driven by FGFR2 fusion proteins. Our work shows that mouse p53-null liver organoids -engineered to express either a sporadic, i.e. F-TACC3, or highly prevalent, i.e. F-BICC1^13^, FGFR2 fusion -generated iCCA lesions with full penetrance upon orthotopic or subcutaneous transplantation in immuno-compromised mice. We have recently extended this work to show that *Tp53*^*-/-*^ mouse liver organoids engineered to express FGFR2-CCDC6 (F-CCDC6)^3^ also yielded iCCA-like lesions upon s.c. transplantation (Supplementary Figure11A-C and Supplementary Table 1).

Besides providing formal demonstration that FFs act as oncogenic drivers in iCCA, our work indicates that structurally different FFs may diverge in terms of oncogenic potential in a p53-null background. Thus, F-MGEA5-expressing organoids were not tumorigenic when transplanted in liver and yielded a *bona fide* iCCA-like lesion in only 1/8 s.c. injection sites. On the other hand, organoids expressing F-CCDC6 generated tumors in 75% of s.c. injection sites (Supplementary Table 1). This gradient of tumorigenic potency correlated with tumor growth kinetics, with F-CCDC6 and F-MGEA5 s.c. lesions developing at a slower pace than F-BICC1 and F-TACC3 tumors (compare Figure 2B with Supplementary Figures 4B and 11B). We note that the quite low tumorigenic activity of F-MGEA5-expressing *Tp53*^*-/-*^ organoids could not be ascribed to a deficit of F-MGEA5 expression/activation in comparison to F-TACC3 and F-BICC1 (Figure 1C). That structural diversity among FFs may impact on their biology was also suggested by the observation that gene expression patterns of F-BICC1- and F-TACC3-driven tumors tended to cluster independently from each other.

The above data point to a degree of, so far unappreciated, heterogeneity among iCCA FFs, at least in a p53-null background. Complementing this observation, a recent study reported that the oncogenic activity of FGFR2-AHCYL1 was genetic context-dependent. Accordingly, FGFR2-AHCYL1-expressing liver organoids became tumorigenic upon loss of *Cdkn2a* expression, but not *Tp53* ablation^9^. Thus, the picture seems to be emerging in which FF-driven iCCA pathogenesis may be orchestrated by an interplay between the genetic background generated by loss of a specific tumor suppressor gene (TSG) on one side and still unclear biological properties intrinsic to individual FFs on the other. To test this model, it will be necessary to compare the oncogenic activity of different FFs upon their expression in mouse liver organoids individually lacking any of the three TSGs that are most often mutated in FF+ iCCA, namely *Tp53, Cdkn2a* and *Bap1*^12, 13^. A prediction stemming from this model is that genotype-specific FF co-dependencies may exist in iCCA. We argue that the above examples illustrate the utility of our iCCA model in generating and testing translationally relevant hypotheses.

Tumoroids and 2D cell lines derived from our iCCA models enabled proof of principle pre-clinical studies that identified Ras-Erk as a necessary effector pathway downstream to FFs. This conclusion was based on different pieces of evidence. First, trametinib, a clinically approved MEK1/2 inhibitor, displayed potent activity in a number of FF+ iCCA cellular models. In line, pharmacological blockade of SHP2, which acts downstream to FFs and upstream of RAS^27^, inhibited Erk activation in iCCA cells and blunted their viability/proliferation. Second, trametinib suppressed the EGF-dependent rescue of BGJ398-mediated growth suppression, implying that Ras-Erk re-activation by parallel Egfr signaling was sufficient to by-pass FF inhibition. Third, dependence on Erk1/2 signal output was manifest also in iCCA cells driven by an FF carrying the V565F mutation, which causes clinical resistance to BGJ398. The notion that Ras-Erk signaling is required downstream to FFs in murine iCCA cells is in line with: a) data pointing to up-regulation of KRAS-driven transcriptional signatures in FF+ human iCCA (Figure 2F); b) KRAS and BRAF acting as oncogenic drivers in human iCCA^1^; c) FGFR2 fusions being mutually exclusive with either *BRAF* or *KRAS* mutations^13^.

Trametinib interacted synergistically with BGJ398 in iCCA cellular models driven by F-BICC1, F-TACC3 (Figure 5A, B) and F-CCDC6 (Supplementary Figure 11D-F), with the B+T combination exerting more potent cell killing activity when compared to single agents. These data converge upon the emerging paradigm that simultaneous blockade of multiple components of an oncogenic signaling axis increases the amplitude of pathway inhibition^31^.

The above *in vitro* results informed pharmacologic studies in mice bearing iCCA-like lesions generated by s.c. transplantation of F-BICC1 tumoroids. Pharmacodynamic experiments confirmed that the B+T combination inhibited Erk activity better than single agent BGJ398 or trametinib. In line, upfront B+T treatment afforded a significantly longer control of tumor growth in comparison to single agent BGJ398. The magnitude of this effect was largest when mice were dosed with BGJ398 and trametinib at 15 mg/Kg and 1 mg/Kg, respectively, for five days per week. Two sudden deaths were recorded in the B+T group at the end of this experiment, possibly raising doubts about cumulative toxicity. We note, however, that two studies had reported no lethal toxicity when the B+T combo was administered to immuno-compromised mice at similar doses and for longer periods of treatment^32, 33^. Scaling down B+T dosage in a second experiment still afforded superior therapeutic efficacy than BGJ398 alone, although control of tumor growth was slightly less durable than that observed with higher B+T dosage. While additional studies will be necessary to determine optimal dosing/scheduling of the B+T combo in our iCCA model, we anticipate that translation of our data to clinical experimentation should be facilitated by the available knowledge about the toxicity profiles of clinically advanced F-TKIs and FDA-approved MEK1/2 inhibitors.

An interesting observation was that B+T mitigated therapeutic resistance in mice that progressed while being treated with single agent BGJ398. In assigning these mice to receive the B+T treatment, we explicitly assumed that resistance to BGJ398 was caused by the activity of collateral pathway/s capable of activating Erk1/2 in the face of FF blockade. This view implied that B+T, rather than single agent trametinib, was bound to be most beneficial. Additional work, which goes beyond the scope of the present study, will be required to formally proof the validity and mechanistic underpinnings of this model.

Our observation that FF+ iCCA cells expressing the FGFR2-TACC3 V565F remained dependent on Ras-Erk signaling suggests that upfront treatment with the B+T combo holds the potential to delay the emergence of resistant clones carrying mutations in the FF TKD^5, 6^. A somewhat similar strategy was recently validated at the preclinical level in tumors driven by NTRK fusions, which were shown to develop resistance to NTRK-specific TKIs via converging mechanisms capable of restoring RAS-ERK signaling^30^.

## Conclusions

We have developed a mouse model that allows for rapid generation of genetically defined FF+ iCCA in immuno-compromised mice. We predict that this model will foster research on the mechanistic bases of FF-driven oncogenic conversion of biliary epithelial cells. We also envision that 2D and 3D cellular models derived from murine iCCA will enable pharmacologic and genetic studies aimed at unbiased discovery of actionable dependencies in FF-driven iCCA cells.

## Supporting information

Supplementary informations

## Acknowledgements

We dedicate our work to the cherished memory of our friends and colleagues Daniela and Marianna, who fought courageously against cancer, and to the Italian Public Health System professionals who operated at the frontline of the Covid-19 epidemic. We are grateful to L. Cantley, H. Clevers, M. Huch, H. Liu, A. Saborowski, E. Salvati, F. Spinella, and our colleagues at the Regina Elena Institute for advice and reagents. We thank S. Alemà and G. Blandino for critical reading of the manuscript.

Author names in bold designate shared co-first authorship.

