## Supplementary informations for "FGFR2 fusion protein-driven mouse models of intrahepatic cholangiocarcinoma unveil a necessary role for Erk signaling"

Short title: *FGFR2 fusions in cholangiocarcinoma models*

Giulia Cristinziano[1], Manuela Porru[2], Dante Lamberti[1], Simonetta Buglioni[3], Francesca Rollo[3], Carla Azzurra Amoreo[3], Isabella Manni[2], Diana Giannarelli[4], Cristina Cristofoletti[5], Giandomenico Russo[5], Mitesh J. Borad[6], Gian Luca Grazi[7], Maria Grazia Diodoro[3], Silvia Giordano[8,9] Mattia Forcato[10], Sergio Anastasi[1]*, Carlo Leonetti[2]* and Oreste Segatto[1]*

[1]Unit of Oncogenomics and Epigenetics, IRCCS Regina Elena National Cancer Institute, Rome, Italy

[2]SAFU, IRCCS Regina Elena National Cancer Institute, Rome, Italy

[3]Department of Pathology, IRCCS Regina Elena National Cancer Institute, Rome, Italy

[4]Unit of Biostatistics, IRCCS Regina Elena National Cancer Institute, Rome, Italy

[5]Istituto Dermopatico dell'Immacolata, IDI-IRCCS, Rome, Italy

[6]Division of Hematology and Oncology, Mayo Clinic, Scottsdale, USA

[7]Division of Hepatobiliary Pancreatic Surgery, IRCCS Regina Elena National Cancer Institute, Rome, Italy

[8] Department of Oncology, University of Torino, Candiolo, Italy

[9] Candiolo Cancer Institute, FPO-IRCCS, Candiolo, Italy

[10]Center for Genome Research, Dept. of Life Sciences, University of Modena and Reggio Emilia, Modena, Italy

*Corresponding Author

**Grant support**: O.S. is funded by AIRC (IG2018, ID 21627, PI Segatto Oreste) and an intramural grant-in-aid funded by the Italian Ministry of Health.

**Please address correspondence to**:

**Oreste Segatto, MD**

Unit of Oncogenomics and Epigenetics,
IRCCS Regina Elena National Cancer Institute,
via E. Chianesi, 53
00144 Rome, Italy
Ph. 39-0652662551


**Carlo Leonetti, PhD**
SAFU
IRCCS Regina Elena National Cancer Institute,
via E. Chianesi, 53
00144 Rome, Italy
Ph. 39-0652662534


**Sergio Anastasi, PhD**
Unit of Oncogenomics and Epigenetics,
IRCCS Regina Elena National Cancer Institute,
via E. Chianesi, 53
00144 Rome, Italy
Ph. 39-0652662568


**All Authors, except M.J.B., have no personal, professional or financial conflicts to disclose.**

**M.J.B. disclosures:** ADC Therapeutics – Consulting to self; Exelixis Pharmaceuticals – Consulting to self; Inspyr Therapeutics – Consulting to self; G1 Therapeutics – Consulting to self; Immunovative Therapies – Consulting to self; OncBioMune Pharmaceuticals – Consulting to self; Western Oncolytics – Consulting to self; Lynx Group – Consulting to self; Genentech – Consulting to self; Merck – Consulting to self; Huya – Consulting to self; Astra Zeneca – Travel Support to self

**Authors' contributions**

M.P., D.L., S.B., G.C.: acquisition of data; analysis and interpretation of data

F.R., C.A.M.: acquisition of data

M.G.D., I.M.: analysis and interpretation of data

D.G.: statistical analysis

M.F.: study concept and design, analysis and interpretation of data, statistical analysis

G.R., M.J.B., G.L.: critical revision of the manuscript for important intellectual content

S.G.: analysis and interpretation of data; drafting of the manuscript, critical revision of the manuscript for important intellectual content

S.A.: acquisition of data; analysis and interpretation of data; drafting of the manuscript, study supervision

C.L.: study concept and design, analysis and interpretation of data, drafting of the manuscript

O.S.: study concept and design, drafting of the manuscript, obtained funding, study supervision

#### Supplementary methods

Lists of reagents and chemicals, antibodies, PCR primers and kits are reported in Supplementary Tables 3-6.

##### Composition of cell culture media

Liver isolation medium is composed by Advanced DMEM/F12 supplemented with 1% (vol/vol) penicillin/streptomycin, 1% (vol/vol) Glutamax, 10 mM HEPES, 1:50 B27 supplement (without vitamin A), 1 mM N-acetyl-l-cysteine, 5% (vol/vol) Rspo-1 conditioned medium, 10 mM nicotinamide, 10 nM recombinant human (Leu15)-gastrin I, 50 ng/ml recombinant mouse EGF, 100 ng/ml recombinant human FGF10, 50 ng/ml recombinant human HGF, 30% (vol/vol) Wnt3a-conditioned medium, 25 ng/ml recombinant human Noggin and 10 µM Rho kinase (ROCK) inhibitor (Y27632), as described[1].

The expansion medium is composed by Advanced DMEM/F12 supplemented with 1% (vol/vol) penicillin/streptomycin, 1% (vol/vol) Glutamax, 10 mM HEPES, 1:50 B27 supplement (without vitamin A), 1 mM N-acetyl-l-cysteine, 5% (vol/vol) Rspo-1 conditioned medium, 10 mM nicotinamide, 10 nM recombinant human (Leu15)-gastrin I, 50 ng/ml recombinant mouse EGF, 100 ng/ml recombinant human FGF10, 50 ng/ml recombinant human HGF, as described[1].

The deprived medium is composed by Advanced DMEM/F12 supplemented with 1% (vol/vol) penicillin/streptomycin, 1% (vol/vol) Glutamax, 10 mM HEPES, B27 supplement at 1:50 dilution (without vitamin A), 1 mM N-acetyl-l-cysteine, 10 mM nicotinamide, and 10 nM recombinant human (Leu15)-gastrin I).

##### Liver organoids

Liver organoids were isolated from adult C57BL/6J *Tp53*$^{-/-}$ mice, which were obtained from Jackson Laboratories, according to published protocols[1]. In detail, murine livers were minced and enzymatically digested for 45-120 minutes at 37 °C in DMEM supplemented with 1% (vol/vol) FBS and containing 0.125 mg/ml of collagenase from *Clostridium histolyticum*, 0.125 mg/ml of dispase II and 0.1 mg/ml DNase I. Ductal structures were collected by hand-picking and centrifuged at 300g for 5 min at 8 °C. The pellet was resuspended in Growth Factor Reduced Matrigel and plated in 24-well plates (50 µl droplet per well). After solidification, Matrigel droplets were overlaid with 500 µl of mouse liver isolation medium. For further passaging, organoids were mechanically disrupted by repeated pipetting and cultured in organoid expansion medium. When required, liquid medium overlaying the matrigel was changed twice a week (0.5-1 ml/well in 24-well plates).

**Generation of 3D and 2D tumor cell cultures**

Tumor-derived organoids (tumoroids) were isolated according to published protocols[2]. Briefly, tumors were minced and enzymatically digested for one to four hours in a shaking incubator at 37 °C in digestion solution (DMEM supplemented with 1% (vol/vol) FBS and 1% (vol/vol) penicillin/streptomycin) with 2.5 mg/ml of collagenase from *Clostridium histolyticum* and 0.1 mg/mL DNase I. Cells were washed, spun at 300g for 5 min at 8 °C and passed through a 100 µm filter. The resulting cell pellet was resuspended in Growth Factor Reduced Matrigel and overlaid with organoid expansion medium. For further passaging, tumoroids were mechanically disrupted by repeated pipetting and cultured in organoid expansion medium. When required, liquid medium overlaying the matrigel was changed twice a week (0.5-1 ml/well in 24-well plates). A fraction of the above tumor-cell pellet was plated in gelatine-coated tissue culture dishes containing DMEM supplemented with 10% (vol/vol) FBS and 1% (vol/vol) penicillin/streptomycin in order to derive 2D cell lines.

**Cell viability assays**

Tumoroid-viability assays were performed according to Broutier *et al.*[2], with minor modifications. Briefly, tumoroids were enzymatically dissociated with TrypLE Express before being suspended in 2% (vol/vol) matrigel-containing deprived medium, dispensed into 96-well plates ($7x10^3$ cells/well) and grown in 2D conditions. Tumor-derived 2D cell lines were plated at $1x10^4$ cells/well into gelatine-coated 96-well plates in DMEM supplemented with 1% (vol/vol) FBS and 1% (vol/vol) penicillin/streptomycin. Assays were run in quadruplicate wells. Cells were allowed to adhere to the plastic overnight (O/N) and then subjected to drug treatment. Cell viability was assayed after 72 hours of drug treatment using the CellTiter-Glo kit (Promega), according to manufacturer's specification. The half maximal inhibitory concentration ($IC_{50}$) was calculated using CalcuSyn Software. Data were computed from at least two independent experiments. Representative photographs of cell cultures were taken at the 72-hour endpoint of growth assays using Incucyte S3 Live-Cell Imaging System (Essen BioScience, Ann Arbor, MI). For combination drug assays, tumoroids were plated in 96-well plates as described above. BGJ398 and trametinib were combined at equipotent concentrations, according to a constant ratio combination of drugs. Cell viability was assessed after 72 hours of drug treatment with the CellTiter-Glo kit. Data from at least three independent experiments were compiled and CalcuSyn Software was used to calculate the Combination Index (CI), according to Chou and Talalay[3]. For long term growth assays, iCCA cell lines were seeded in gelatin-coated 6-well plates ($6x10^4$/well) in DMEM supplemented with 5% (vol/vol) FBS, glutamax and 1% (vol/vol) penicillin/streptomycin. Drug treatments were started the next day. Assays were stopped when untreated control cultures reached confluence, which usually took about a week. Plates were fixed with 4% (vol/vol) paraformaldehyde (PFA) and then stained with 0.5% (w/vol) crystal violet. Data were obtained from at least three independent experiments.

**Lentiviral and retroviral vectors**

Previously described cDNAs encoding epitope-tagged FGFR2-BICC1, FGFR2-MGEA5, FGFR2-TACC3 and FGFR2-TACC3 V565F[4] were subcloned into the pMSCV Puro retroviral vector, which provides for optimal expression in stem cells. The pBabepuro-FGFR2-CCDC6 retroviral vector was kindly provided by Dr. L. Cantley. Note that FGFR2IIIb residues are numbered in this paper according to sequence NM_001144913.1. For the sake of clarity, we note that previous studies[4-6] reported numbering of FGFR2 residues in iCCA fusions according to the FGFR2IIIc reference sequence NM_000141.4. pRRLSIN.cPPT.RFPL4b.Luciferase.WPRE was a gift from Stephen Tapscott (Addgene plasmid # 69252; http://n2t.net/addgene:69252; RRID:Addgene_69252).

**Generation of recombinant lentivirus and retrovirus**

293FT cells (Thermo Fisher Scientific) were seeded at $2.5x10^6$ in 60 mm dishes. The following day, cells were transfected with Lipofectamine 3000 (Thermo Fisher Scientific) according to the manufacturer's specification, using 10.4 µg of plasmid DNA per each dish (5.6, 3.0 and 1.8 µg of transfer, psPAX2 packaging and pMD2.G envelope plasmid DNA, respectively, which corresponds to a 3.1:1.7:1 molar ratio). Transfection medium was removed after 6 hours and replaced with 5 ml/ dish of DMEM containing 10% (vol/vol) FBS. Virus-containing medium was harvested 24 and 48 hours post transfection. After centrifugation, the supernatant-containing virus was filtered through a 0,45 µm filter to remove cell debris, aliquoted and stored at -80°C.

The ecotropic Phoenix packaging cell line[7] was used to generate stocks of replication-defective recombinant retrovirus. Cells were seeded at $1.8x10^7$ in 150 mm dishes. The next day, cells were transfected with Lipofectamine 3000 according to the manufacturer's specification, using 70 µg of transfer plasmid DNA per dish. Transfection was terminated after 6 hours by changing medium. Virus-containing medium was harvested 24 and 48 hours past transfection. Virus was concentrated by polyethylene glycole (PEG) precipitation using a kit manufactured by Abcam. PEG-precipitated virus particles were collected by centrifugation and resuspended in organoid transduction medium (isolation medium supplemented with 8 µg/ml polybrene) to obtain a final 100X concentration, aliquoted and stored at -80°C.

**Infection procedures**

Organoids infection was carried out according to a published protocol[8] with some modifications. In brief, organoids were enzymatically dissociated with TrypLE Express and seeded into 24-well plates ($5.0x10^4$ cells/well) in 2D culture conditions using expansion medium containing 2% (vol/vol) matrigel. The next day, the medium was replaced with concentrated retrovirus stock diluted to a final 60X concentration (corresponding to a nominal 5-10 multiplicity of infection value). Infection was carried out for 6-8 hours at 32 °C in 5% $CO_2$ atmosphere and terminated by changing medium. The next day, organoids were super-infected with a luciferase-encoding lentivirus (pRRLSIN.cPPTLuciferase.WPRE from Addgene) for 6-8 hours at 37 °C in 5% $CO_2$ atmosphere. Puromycin selection (1 µg/ml) was started 2-3 days past retrovirus infection. After 72 hours, a time interval sufficient to obtain >90% cell death in control mock-infected cultures, puromycin-selected cells were transferred to 3D culture conditions and kept under puromycin selection for at least two more weeks.

**RNA extraction and RNA sequencing**

Total RNA was extracted from mouse normal livers (n = 2) and murine iCCA tumors (n = 12, of which n = 3 F-BICC1 i.h. tumors, n = 2 F-BICC1 s.c. tumors, n = 3 F-TACC3 i.h. tumors, n = 2 F-TACC3 s.c. tumors and n = 2 s.c. F-TACC3 obtained upon transplantation of F-TACC3 tumoroids) with QIAzol and purified with the miRNeasy Mini kit (Qiagen). RNA was quantified using the Qubit® RNA HS Assay Kit (Thermo Fischer Scientific). RNA integrity (RIN) was measured using the Bioanalyzer 2100 RNA 6000 Nano Kit (Agilent Technologies). High-quality total RNA was assessed for all samples, with RIN values ranging from 8.0 to 10.0. TruSeq® Stranded mRNA Kit (Illumina) was used to generate all libraries, according to the manufacturer's recommendations. Produced libraries were quantified by Qubit® DNA HS Assay Kit (Thermo Fischer Scientific) and Bioanalyzer High Sensitivity DNA Kit (Agilent Technologies). An equimolar libraries pool (14 samples) was loaded onto Illumina NextSeq 500 platform into a High Output Kit v2.5 cartridge (2x75 bp paired-end sequencing with about 25 million clusters per sample). Base-calling was performed by Illumina Real-Time Analysis (RTA) software and NextSeq Control Software (NCS).

**Analysis of RNA-seq data**

Raw reads from murine iCCA and normal liver were aligned using STAR[9] version 2.5.3a to build version mm10 of the mouse genome. Counts for UCSC annotated genes were calculated from the aligned reads using *featureCounts* function of the Rsubread R package[10]. Normalization was carried out using edgeR package[11] and R (version 3.3.1). Raw counts were normalized to obtain Counts Per Million mapped reads (CPM). Only genes with a CPM greater than 1 in at least 1 sample were retained for downstream analysis. Principal component analysis was performed on log normalized data using the most variable 10% of genes. RNA-seq data from this study have been deposited in the Gene Expression Omnibus database under accession number GSE150504.

Raw counts for TCGA cholangiocarcinoma samples were downloaded from TCGA CHOL project[12] using TCGAbiolinks R package[13]. We discarded samples with “extrahepatic biliar duct” or “gallbladder” tissue of origin, with HBV positive status or atypical FGFR2-FRK fusion; to exclude samples with an IDH mutant signature, we focused on samples belonging to mRNA-seq cluster 3, for a total of 4 FF+ and 11 FF- samples. Normalization was carried out as for murine samples. Gene set enrichment analysis (GSEA) was performed on TMM normalized data and the Hallmark collection of Molecular Signature Database (MSigDB[14]) version 7.0. Single sample gene set variation analysis (GSVA) was performed on iCCA samples of the murine and TCGA data with GSVA package[15] and the MSigDb Hallmark collection version 7.0.

**RT-PCR studies**

Organoids were resuspended in 1 ml of TRIzol. RNA was extracted and purified according to manufacturer’s instructions and dissolved in RNAse-free $H_2O$. Complementary DNA (cDNA) was synthesized using EuroRT M-MLV Reverse Transcriptase and Random Hexamers. cDNA was subjected to RT-PCR using 0.4 μM forward and reverse primers and Wonder Taq Hot Start Thermostable DNA polymerase in a 25 μl final volume reaction. Amplification was carried out in a TPersonal Thermocycler (Biometra). Conditions for RT PCR were: 1 min at 95 °C for the initial denaturation and 30 cycles at 95 °C for 30 sec, 58 °C for 1 min and 72 °C for 1 min. For Real-Time analyses forward and reverse primers were used each at 0.3 μM and 10 μl of 2X Sybr Green Master Mix was added to a final reaction volume of 20 μl. Levels of cDNA were normalized to the *Actb* control housekeeping gene. Reactions were run in duplicate in a StepOne™ Real-Time PCR System (Applied Biosystems). At the end of the assay, a melting curve analysis of dissociation of double strands during heating was performed as a quality control. Relative gene expression was evaluated using the $2^{-\Delta\Delta CT}$ method. Real-time PCR assays were repeated twice to derive mean values ±

standard error of the mean (SEM). Primers used in RT and Real-Time PCR experiments are listed in Supplementary Table 5.

**Immunoblot analyses**

Cells were lysed on ice in RIPA buffer (50 mm Tris-HCl pH 7.5, 150 mM NaCl, 1 mM EDTA, 1% (vol/vol) Triton X-100, 0.2% (w/vol) sodium deoxycholate and 0.1% (w/vol) SDS) containing a cocktail of protease inhibitors and 1 mM sodium vanadate. Lysates were cleared by centrifugation. Protein concentration was determined by the Pierce BCA protein assay kit (Thermo Fisher Scientific). Proteins were resolved in SDS-PAGE and transferred to nitrocellulose membranes. Membranes were blocked in TBS (25 mM Tris-base, 3 mM KCl, 140 mM NaCl, pH 7.4) containing 0.05% (vol/vol) Tween-20 and 5% (w/vol) BSA and incubated O/N at 4 °C with primary antibody. Primary antibodies are listed in Supplementary Table 4. The anti-MYC 9E10 mouse hybridoma was purified in house and used at 2 μg/ml. After incubation with primary antibody, membranes were washed in TBS and incubated for 1 h at 22 °C with HRP-conjugated secondary antibodies. Immunoreactivity was detected by ECL (GE Healthcare) as suggested by the manufacturer. Immunoreactivity was imaged using a Uvitec chemiluminescence analyser.

**GST pulldown assays**

GEX-based vectors for bacterial expression of GST and GST-GRB2 SH2 were described[16]. pGEX SHP-2(NC)-SH2[17] was a gift from Bruce Mayer (Addgene plasmid #46499; http://n2t.net/addgene:46499; RRID:Addgene_46499). Recombinant GST proteins were purified from bacteria lysates and immobilized onto glutathione-agarose beads (GE Healthcare) as described[16]. Total cell lysates were made in HNTG buffer (50 mM HEPES pH 7.5, 150 mM NaCl, 10% (vol/vol) glycerol, 1% (vol/vol) Triton X-100, 5 mM EGTA) containing protease inhibitors and 1 mM sodium vanadate. Clarified cell lysates (500 μg per assay) were incubated for 2.5 hr at 4 °C under rotation with glutathione agarose beads bound to about 10 μg GST-fusion protein per assay. At the end of incubation time, beads were washed and protein complexes analysed by immunoblotting. Blot sections containing GST-fusion proteins were stained with Ponceau's red.

**Immunofluorescence**

Organoids were plated onto 8-well chamber-slides ($1x10^4$ cells/small cell clumps per each well). Tumor-derived 2D cell lines were plated in 35 mm gelatin-coated dishes ($3x10^5$ cells/dish). 3D and 2D cultures were fixed with 4% (vol/vol) PFA (diluted in PBS from 37% stock) for 20 min and subsequently permeabilized with 0.5% (vol/vol) Triton X-100 in PBS for 20 min at 22 °C. Cells were incubated with 1% (w/vol) BSA in PBS containing 0.2% (vol/vol) Triton X-100 and 0.05% (vol/vol) Tween-20 for 1 hour to block non-specific immuno-reactivity and then incubated with primary antibodies at 4 °C O/N (see Supplementary Table 4). The following day, cells were incubated in the dark with secondary antibodies for 1 hour at room temperature. When required, this step was followed by incubation with phalloidin-tetramethylrhodamine B isothiocyanate (TRITC) for 30 min at room temperature to visualize F-actin. Nuclei were stained with DAPI. Fluorescence was imaged using an Olympus BX53 microscope with epifluorescence; photographs were taken (×20 and x40 objectives) with a cooled camera device (ProgRes MF).

**Drug preparations for animal studies**

BGJ398 was dissolved in 30% (vol/vol) PEG-400, 0.5% (vol/vol) Tween-80, 5% (vol/vol) polypropylene glycol. Trametinib was dissolved in an aqueous mixture of 0.5% (w/vol) hydroxypropyl methyl cellulose (HPMC) and 0.2% (vol/vol) Tween-80.

**Mouse genotyping**

Tail biopsies of C57BL/6J mice were digested O/N at 56 °C with a solution containing 0.1 mg/ml Proteinase K, 0.45% (vol/vol) NP-40, 0.45% (vol/vol) Tween-20, 0.01% (w/vol) gelatin, 50 mM KCl, 10 mM Tris HCl pH 7.5, 1.5 mM $MgCl_2$. The next day, Proteinase K was inactivated at 95 °C for 5 min. Samples were centrifuged at 4 °C for 5 min and were immediately used for PCR. Genomic DNA was PCR-amplified with 0.4 μM primers using Wonder Taq Hot Start Thermostable DNA polymerase in a 25 μl final volume reaction. The following primers were used:

p53-X6_F: 5'-AGC GTG GTG GTA CCT TAT GAG C-3'

p53-Neo19_F: 5'-GCT ATC AGG ACATAG CGT TGG C-3'

p53-X7_R: 5’-GGA TGG TGG TAT ACT CAG AGC C-3’

Amplification was carried out in a TPersonal Thermocycler (Biometra). Conditions for PCR were: 2 min at 95 °C for the initial denaturation and 35 cycles at 95 °C for 1 min, 62 °C for 1 min and 72 °C for 1.30 min. Expected amplicon size was as follows: 457 bp for wild type, 587 bp for KO (Neo).

**Surgical procedures in mice**

Mice were anesthetized with a combination of tiletamine–zolazepam (Telazol, Virbac, Carros, France) and xylazine (xylazine/Rompun BAYER) given intramuscularly at 2 mg/kg. For intrahepatic tumorigenicity assays, surgery was conducted on liver left lobe along the left rib edge of anesthetized mice. Organoids ($5x10^5$ cells per animal) were suspended in 25 µl matrigel and injected with a syringe using a 28-gauge needle. The injection site was gently pressed with cotton to reduce bleeding and leakage of cells from the injection site. Peritoneum and skin were closed with sutures. For intravital imaging, anesthetized mice were injected intra-peritoneally with 150 mg/kg D-luciferin and imaged once a week following tumoroid transplantation.

For subcutaneous tumorigenicity assays, organoids ($5x10^5$ cells per animal) were suspended in 100 µl volume (50 µl PBS + 50 µl matrigel) and injected into flanks. Subcutaneous tumor growth was monitored by caliper measurements three times per week and tumor volume calculated with the following formula: volume = ($width^2$ x length)/2. When required, mice were anesthetized and euthanized by cervical dislocation following anesthesia.

**Histopathology and immunohistochemistry**

Tumors were removed immediately after sacrifice, fixed in 4% (vol/vol) buffered formalin and paraffin-embedded. Sections were cut 5 µm thick. Hematoxylin and eosin staining was performed according to routine histological practice. Immunohistochemistry analyses were conducted using the following primary antibodies (see Supplementary Table 4 for details): anti-Ck19, anti-FGFR2, anti-phospho-p44/42 MAPK (ERK1/2) and anti-MYC 9E10 mouse hybridoma - all of them after heat induced antigen retrieval in sodium citrate buffer at pH 6.0; anti-HepPar1 - after heat induced antigen retrieval in Tris EDTA buffer at pH 9.0. Cycling cells were stained with anti-Ki67 antibody

- after heat induced antigen retrieval in sodium citrate buffer at pH 6.0. Stromal cells were visualized by Masson's trichrome stain, according to the kit's specification. Immunoreactivity was revealed by Bond polymer Refine Detection, a biotin-free, polymeric horseradish peroxidase (HRP)-linker antibody conjugate system in an automated autostainer (Bond™-III, Leica Biosystem, Milan, Italy), according to the manufacturer's instructions.

Author names in bold designate shared co-first authorship.

#### Legend to Supplementary Figures

**Supplementary Figure 1.** Schematic structure of FGFR2 and representative FGFR2 fusions, i.e. FGFR2-BICC1 (F-B), FGFR2-MGEA5 (F-M) and FGFR2-TACC3 (F-T). FGFR2 fusions share a common N-terminal portion spanning FGFR2IIIb residues 1-768 fused in frame to sequences encoded by fusion genes. Note that the breakpoint leaves an intact FGFR2 tyrosine kinase domain (TKD) in all fusions. Protein-protein interaction motifs/domain such as coiled coil sequences (CC) and sterile alpha motif (SAM) present in fusion sequences are indicated. KH indicates K homology domains present in BICC1.

**Supplementary Figure 2.** (*A*) PCR genotyping of 1-month-old C57BL/6J Tp53 mice. PCR products were separated in a 2% agarose gel. The lower band represents the wild-type band (457 bp), the upper band represents the KO (587 bp) band. Both DNA fragments are present in genomic DNA from heterozygous mice. Molecular weight markers were loaded in the leftmost lane. $H_2O$ was used as negative control. (*B*) Enlarged view of representative immunofluorescence images of FF-expressing organoids, as shown in Figure 1B. Images highlight plasmamembrane localization of the indicated FGFR2 fusion proteins. Scale bar: 100 µm.

**Supplementary Figure 3.** Oncogenic activity of mouse liver organoids expressing F-TACC3. (*A*) Representative images of liver (left) and subcutaneous (right) tumors obtained upon transplantation of F-TACC3 liver organoids. (*B, C*) Histopathology and IHC analysis of representative liver (B) and s.c. (C) tumors obtained upon transplantation of liver organoids expressing F-TACC3. Tissue slides were stained as indicated. The red dotted lines demarcate the boundary between normal liver and tumor. Note marked anti-Ki67 reactivity in tumor areas. (*D*) Principal Component Analysis (PCA) on log normalized counts from murine RNA-seq data. Samples are colored according to their origin (normal liver, murine iCCA driven by F-BICC1 or F-TACC3). Dot shape distinguishes tumors occurring after intra-hepatic (i.h.) or subcutaneous (s.c) organoid transplantation. Two tumors were obtained after s.c. transplantation of F-TACC3 tumoroids and are designed as second-generation tumors (2nd gen). Note that iCCA samples are clearly separated from liver samples. F-BICC1 and F-TACC3 samples tend to cluster independently from each other, with a single exception in each subgroup, regardless of i.h or s.c. origin.

**Supplementary Figure 4.** Tumorigenicity of *Tp53*$^{-/-}$ liver organoids expressing F-MGEA5. (*A*) Representative *in vivo* imaging of orthotopic transplants of F-MGEA5 expressing cells. F-MGEA5 expression/activation in liver organoids is documented in Figure 1C. (*B*) The plot documents the time-dependent volume increase of the single iCCA-like tumor obtained upon s.c. transplantation of F-MGEA5 organoids. Note the slower growth kinetics in comparison to F-BICC1 and F-TACC3 s.c. tumors presented in Figure 2B. (*C*) Gross (left) and microscopic pathology (right) of F-MGEA expressing s.c. iCCA-like tumor. The H&E stain (200x magnification) documents a typical adenocarcinoma histology. (*D*) Top left: macroscopic appearance of a representative cystic lesion, selected among those that developed in 7/8 s.c transplantation sites of F-MGEA5 organoids. H&E stain revealed prevalent aspects of cystadenoma with cystic cavities of variable diameter, lined by a dysplastic epithelium (bottom row, 50x magnification). In an isolated case, a field showing clear-cut progression (red arrow) to cystic adenocarcinoma with micro-papillary projections in the cystic lumen was observed (top row, right, 100x magnification). (*E*) A tumoroid was derived from the F-MGEA iCCA-like tumor described in panel C. Total cell lysates from this tumoroid were immunoblotted with the indicated antibodies along with control lysates from Tum 6 tumoroid (F-TACC3, see Figure 3A). Note that F-MGEA5 expression/activation is comparable to that of F-TACC3. The dotted red line indicates removal of non-relevant intervening lanes from the autoradiogram. Uniform background documents that images of both lanes were taken from the same autoradiogram.

**Supplementary Figure 5.** (*A*) CL 1 and CL 4 iCCA cell lines express the same FF as their tumor of origin, as assessed by MYC immunostaining (F-BICC1 in CL 1) and GFP imaging (F-TACC3 in CL 4). Both cell lines express the cholangiocyte marker Ck19 (red). Scale bar: 50 μm. (*B*) Total cell lysates from the indicated cell lines were immunoblotted with the indicated antibodies. Note the higher stoichiometry of Tyr phosphorylation in F-BICC1 compared to F-TACC3. The dotted red line indicates removal of non-relevant intervening lanes from the autoradiography. Uniform background documents that images of both lanes were taken from the same autoradiogram. (*C*) Representative micrographs of CL 1 (top) and CL 4 (bottom) cells treated for 72 hours with the indicated concentrations of BGJ398. Micrographs were taken with Incucyte S3 Live-Cell Imaging System and relate to growth assays reported in Figure 3D, E.

**Supplementary Figure 6.** Characterization of downstream signaling in FF-expressing iCCA cell lines. (*A*) The indicated F-BICC1 and F-TACC3 iCCA cell lines were treated for 6 or 30 hours with the indicated concentration of BGJ398 before lysis. Total cell lysates were immunoblotted with the indicated antibodies, followed by ECL detection. Results relate to those presented in Figure 4A. (*B*, *C*) Total cell lysates from control and BGJ398-treated CL 1 cells (F-BICC1) were incubated for 2 hours with agarose beads conjugated to bacterially expressed GST, GST-GRB2 SH2 and GST-SHP2 (N+C) SH2. Protein complexes were recovered from beads by boiling in Laemmli buffer and immunoblotted with anti-pFrs2α (B) and anti-pShp2 (C) antibodies. Note that pTyr-SH2 interactions were abolished by BGJ398 treatment.

**Supplementary Figure 7.** Role of Shp2 and Mek1/2 in shaping oncogenic addiction to FF signaling in murine iCCA cellular models. (*A, B*) The indicated cell lines (F-BICC1-expressing CL 1 and CL 2 in panel A; F-TACC3-expressing CL 3 and CL 4 in panel B) were incubated for 72 hours with escalating doses of trametinib. Cell growth was determined with the CellTiter-Glo assay kit. $IC_{50}$ values are reported. (*C, D*) F-BICC1-expressing Tum 1 (C) and CL 1 (D) cells were exposed to escalating doses of SHP099 for 72 hours. Cell growth was determined with the CellTiter-Glo assay kit. (*E*) This panel reports the same dataset described in Figure 4E, adopting a complementary analysis of the data. Drug vehicle-treated controls were made equal to 100, whether EGF-treated (black bars) or not (white bars). In the -EGF series, growth of BGJ398-treated cells was 18.4% of the respective drug-free control. In the +EGF series (black bars), growth of BGJ398-treated cells was 62.3% of the respective drug-free control, implying that EGF was capable of largely rescuing loss of growth/viability imposed by BGJ398 treatment. Note that EGF treatment was inconsequential whenever cells were treated with trametinib, consistent with EGF-dependent rescue being enforced via Ras-Erk reactivation in BGJ398-treated cells.

**Supplementary Figure 8.** Analysis of iCCA lesions obtained upon transplantation of *Tp53*$^{-/-}$ liver organoids expressing FGFR2-TACC3 V565F and evaluation of F-TACC3 V565F dependence on Ras-Erk activation. (*A, B*) Time-resolved analysis of the growth of malignant lesions obtained upon intrahepatic (A) or s.c. (B) transplantation of *Tp53*$^{-/-}$ liver organoids expressing F-TACC3 V565F (see Figure 1C for expression/activation levels of F-TACC3 V565F organoids). (*C*) Representative photographs of liver (42 days post transplantation) and s.c. (35 days post transplantation) F-TACC3 V565F tumors. (*D*) Representative histopathological examination of intrahepatic (i.h., top row) and subcutaneous (s.c., bottom row) tumors expressing F-TACC3 V565F. Note that F-TACC3 V565F is tagged with a 6x MYC epitope. (*E*) Cell lysates from the F-TACC3 V565F-expressing Tum 7 and Tum 8 tumoroids were probed with the indicated antibodies to demonstrate FF expression/activation. (*F*) Tum 7 and Tum 8 cells were incubated for 72 hours with escalating doses of BGJ398. Cell growth was determined with the CellTiter-Glo assay kit. Note lack of cell growth inhibition by BGJ398 at doses at least 10-fold higher than $IC_{50}$ values registered for F-TACC3 tumoroids (Figure 3C) (*G*) Tum 7 and Tum 8 cells were incubated for 72 hours with the indicated doses of trametinib. Cell growth was determined with the CellTiter-Glo assay kit.

**Supplementary Figure 9.** Lack of specific reactivity of anti-pFGFR antibody (Abcam 59194) against FGFR2 fusions. (*A*, *B*) CL 1 cells, which express the F-BICC1 fusion, were treated with either carrier or BGJ398 (100 nM for 60 min) before being processed for IHC analysis (after fixation and paraffin embedding) (A) or cell lysis and WB analysis (B) with the indicated antibodies. Panel A shows prevalent membrane localization of F-BICC1, as visualized by anti-FGFR2 antibodies, a pattern which is not altered by BGJ398 treatment, as expected. The reactivity of Ab 59194 is localized in the cytoplasm and is not altered by BGJ398 treatment. Panel B shows that anti-pFGFR antibody 3471 (Cell Signaling Technology, suitable for WB analysis, but not IHC) identifies Tyr-phosphorylated F-BICC1. As expected, this immunoreactivity is abolished by BGJ398.

**Supplementary Figure 10.** This figure is related to data presented in Figure 6 and Figure 7. (*A*) Body weight values of mice grouped as described in Figure 6A, B were recorded every other day and are reported for each group as average changes from the start of treatment. (*B*) Waterfall plot showing tumor volume changes from baseline as recorded on day 15 for each of the mice belonging to the BGJ398 (dark red bars) and B+T (dark blue bars) groups in Figure 7A. **Dark red bars:** the vertical dotted line separates mice allocated to the B/B+T-1 (left) and B/B+T-2 (right) subgroups subjected to further therapeutic treatment (see yellow shaded area in Figure 7D). **Dark blue bars:** the five leftmost mice, all showing tumor volume reduction >70% (the -70% limit is indicated by the horizontal dotted line), were assigned to the B+T-1 subgroup, while the five rightmost mice were assigned to the B+T-2 subgroup (see Figure 7D, E). (*C*) Body weight values of mice grouped as described in Figure 7D were recorded every other day and reported as average changes from the start of treatment. Please focus on average body weight values of mice in the B+T group, subsequently divided into the B+T-1 and B+T-2 subgoups: note that mice in B+T-1 subgroup showed an average weight loss value of 8.5% on day 12 (end of second cycle of therapy), but regained weight upon drug discontinuation in the third week of the experiment. (*D, E*) The spaghetti plots indicate tumor volume changes in each of the mice initially assigned to the BGJ398 group (see Figure 7D) and subsequently randomized to the B/B+T-1 (D) and B/B+T-2 (E) subgroups. As reference, the spaghetti plot relative to the BGJ398 group of experiment #1 (see Figure 6B) is reported in each panel (black dotted lines).

**Supplementary Figure 11.** *Tp53*$^{-/-}$ liver organoids expressing F-CCDC6 are tumorigenic and generate iCCA-like tumors addicted to F-CCDC6 signaling via Ras-Erk. (*A*) Biochemical analysis of F-CCDC6-expressing (F-C) liver organoids. Total cell lysates were probed with the indicated antibodies. Total cell lysate from an F-TACC3 organoid (F-T) was analyzed as reference. Note the difference in electrophoretic mobility of F-C and F-T, which reflects their different MW. The dotted red line indicates removal of non-relevant intervening lanes from the autoradiogram. Uniform background documents that images of both lanes were taken from the same autoradiogram. (*B*) Time-resolved analysis of s.c tumor growth in mice transplanted with F-CCDC6 liver organoids. Tumors were obtained in 6 out of 8 injection sites. Each time point reports mean volume values ± SEM for these tumors. Note slower growth kinetics in comparison to s.c. F-BICC1 and F-TACC3 tumors (Figure 2B). (*C*) Gross pathology (top) and H&E stain (100x magnification) of representative tissue slides from s.c. F-CCDC6 tumors, including adenocarcinoma with well-developed ductal structures (middle) and carcinoma lacking defined glandular aspects (bottom). Scale bar: 30 μm. (*D*, *E*) CellTiter-Glo assays (72 hours) showing dose-dependent growth suppression of F-CCDC6-expressing tumoroids by BGJ398 (D) and trametinib (E). (*F*) Combination index calculated according to Chou-Talalay for the B+T combo in Tum 10 F-CCDC6 tumoroids. CI <1 indicates synergistic drug interactions.

### Supplementary Figure 1

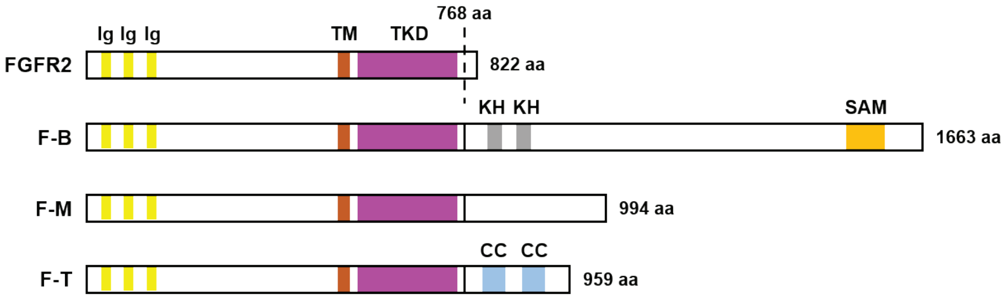

### Supplementary Figure 2

A

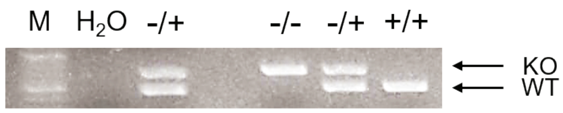


B

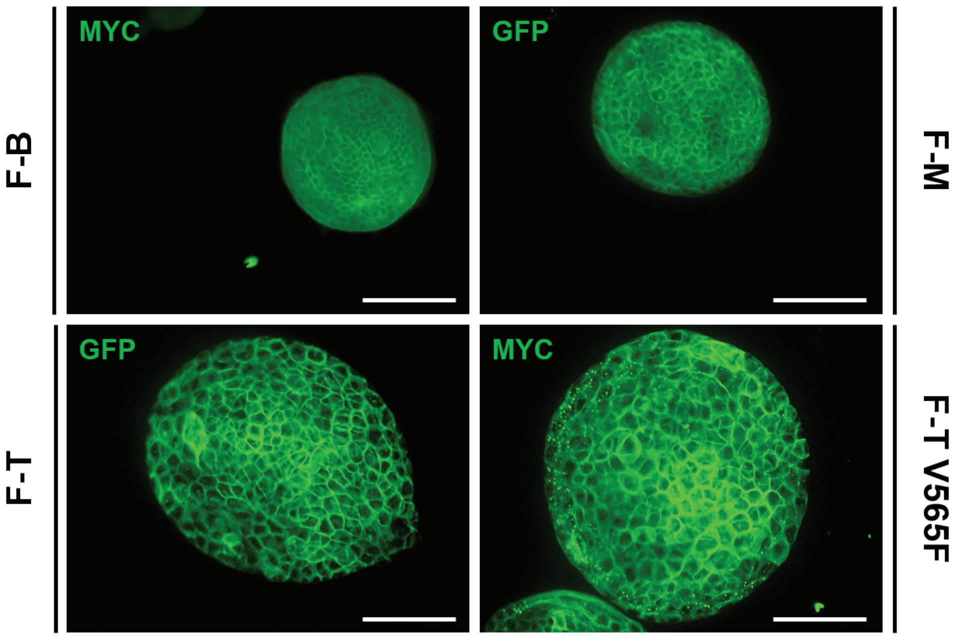

### Supplementary Figure 3

## A

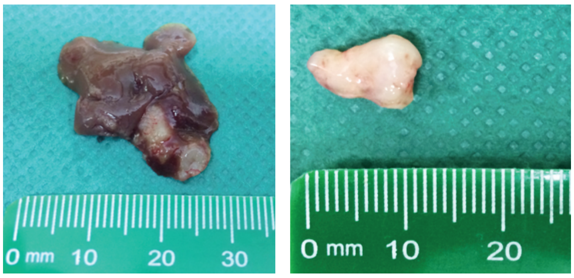


## B

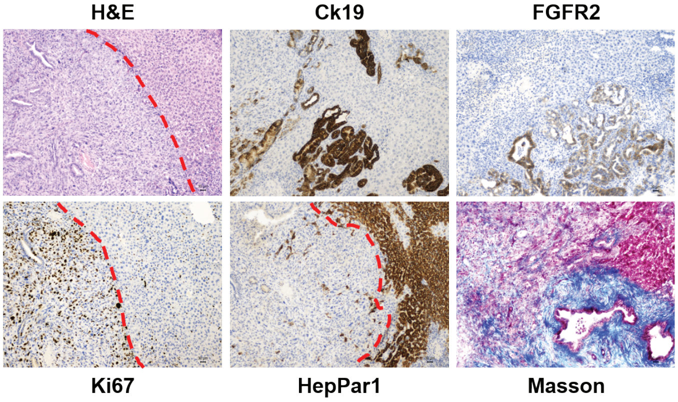


## C

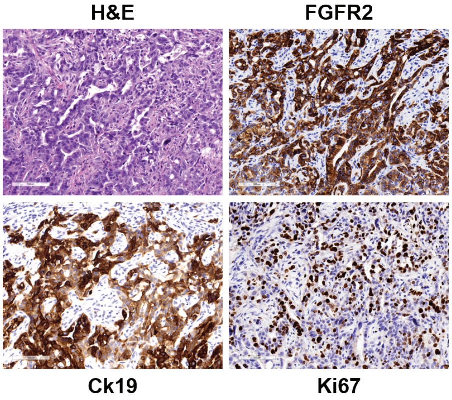


## D

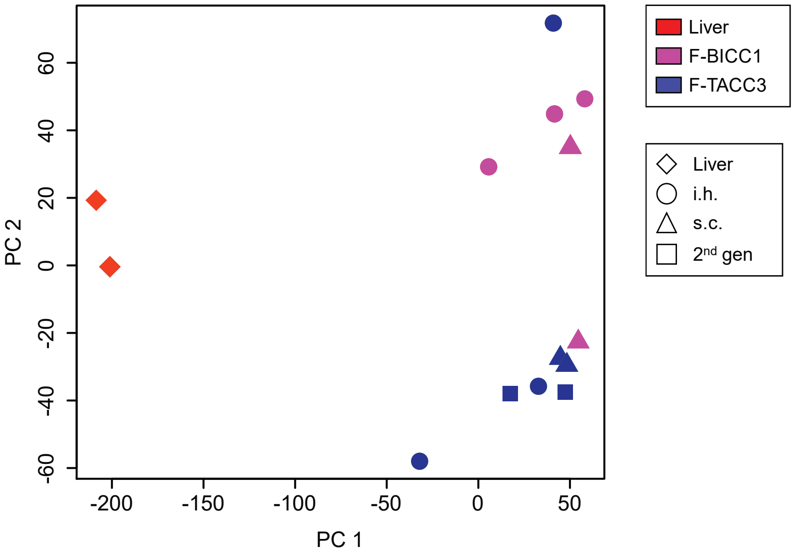

### Supplementary Figure 4

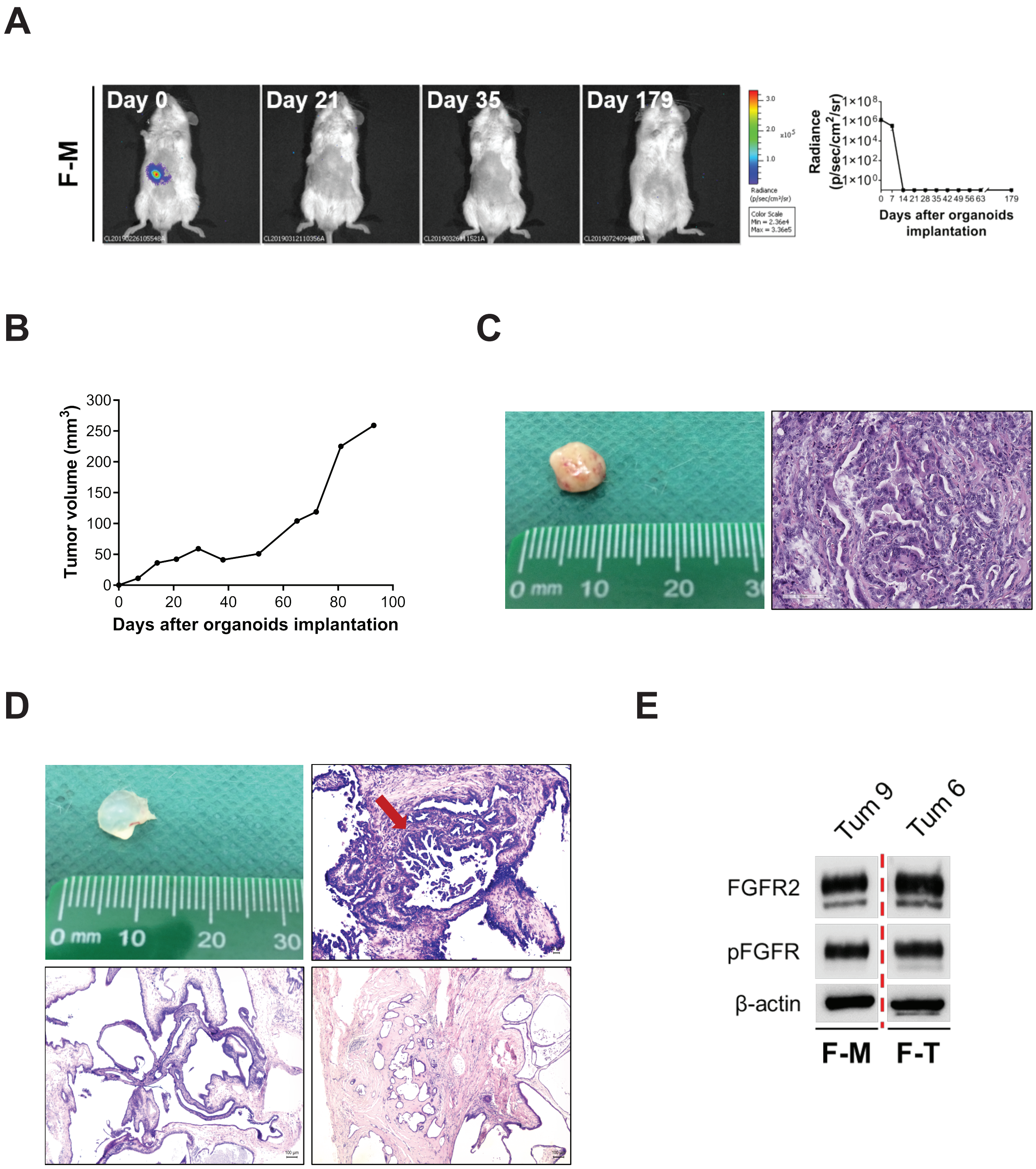

### Supplementary Figure 5

A

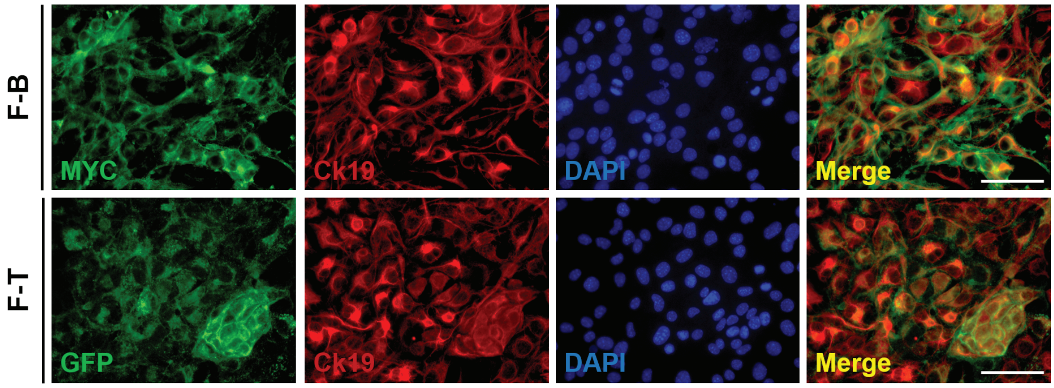


B

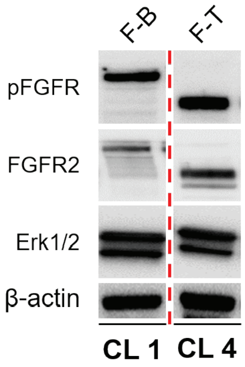


C

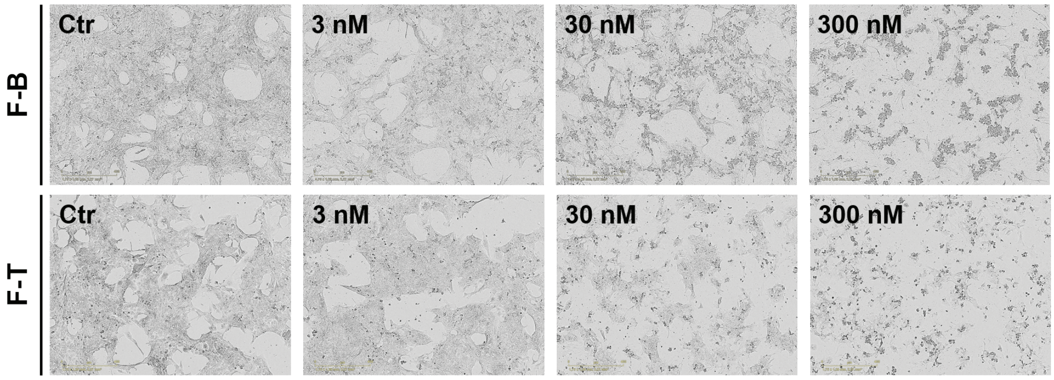

### Supplementary Figure 6

A

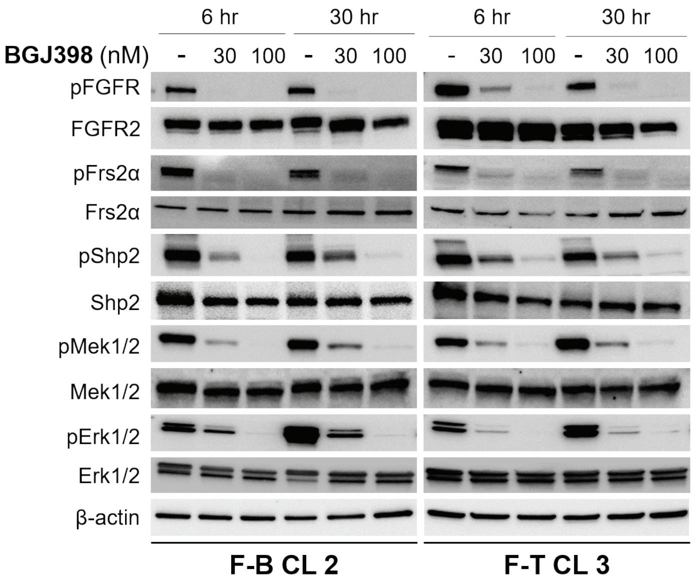


B

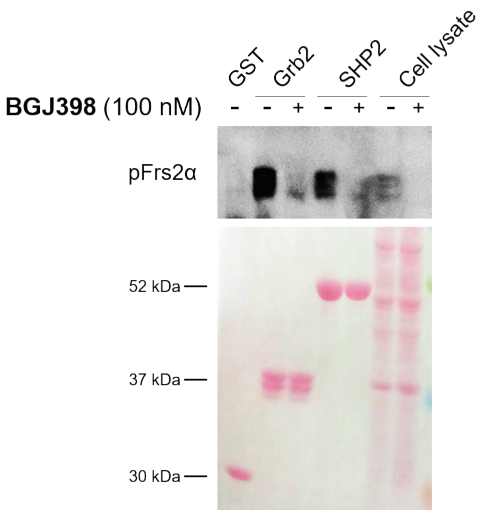


C

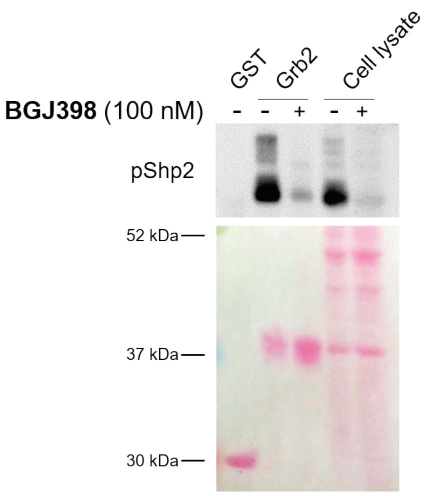

### Supplementary Figure 7

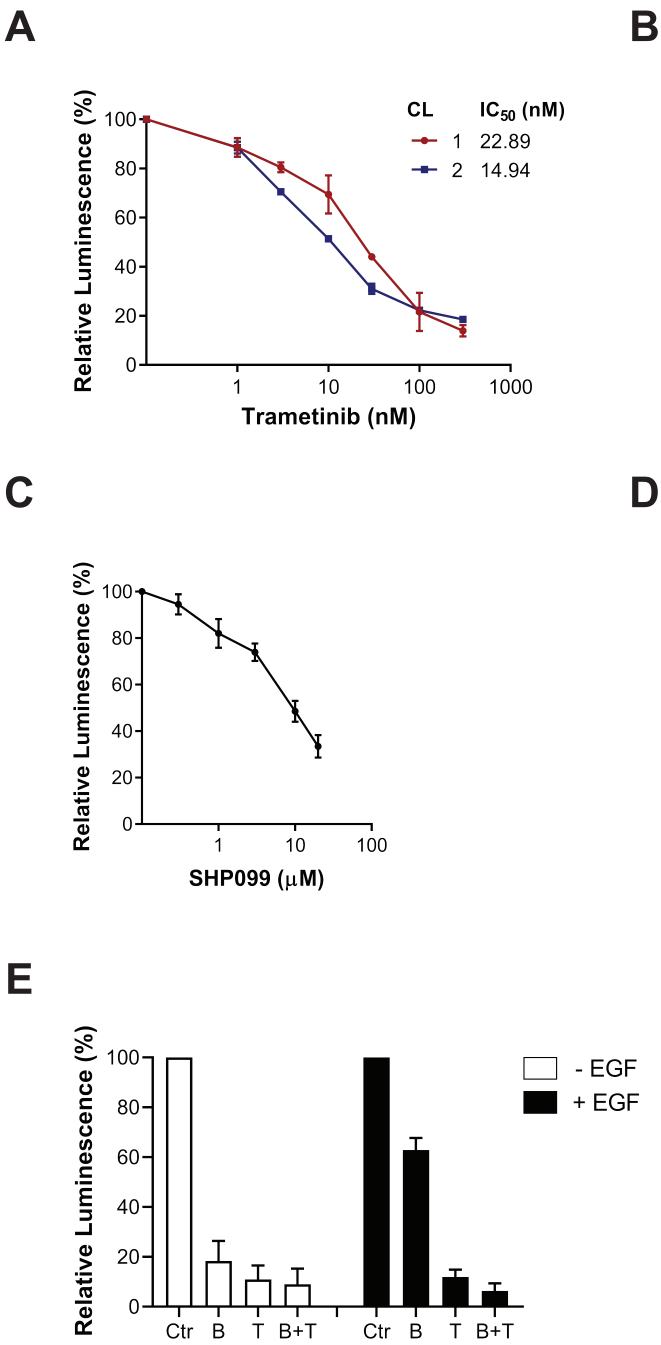

### Supplementary Figure 8

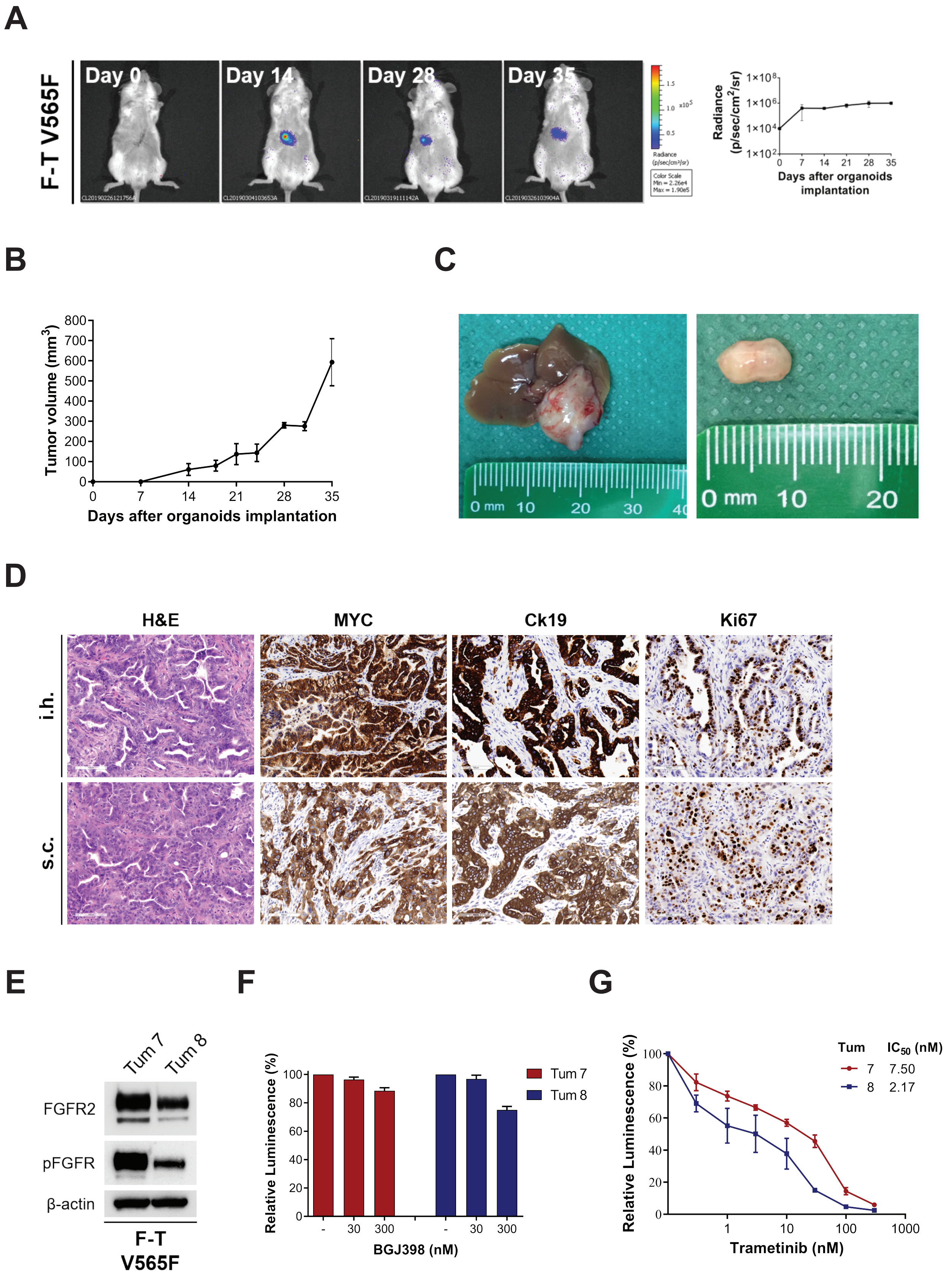

#### Supplementary Figure 9

A

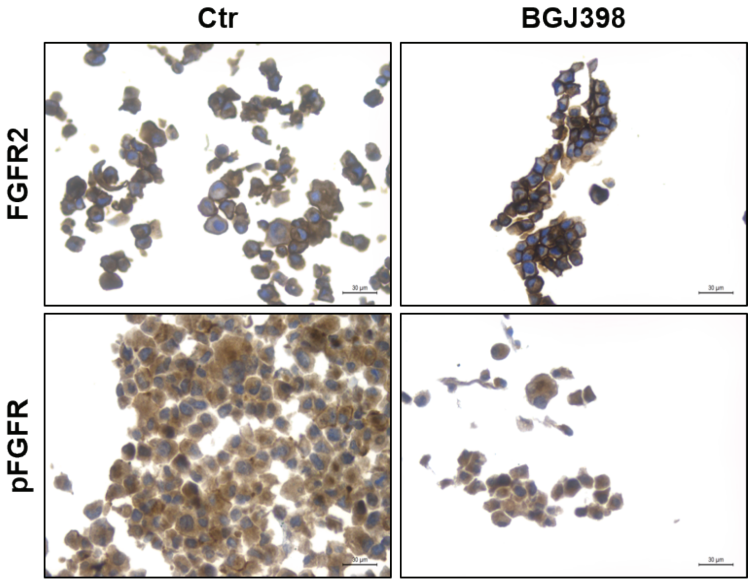


B

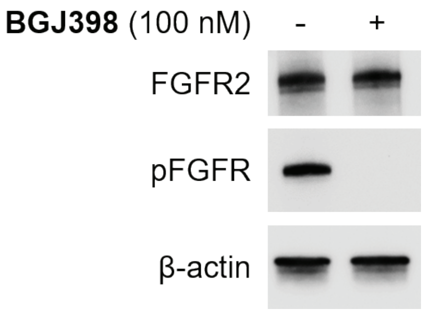

### Supplementary Figure 10

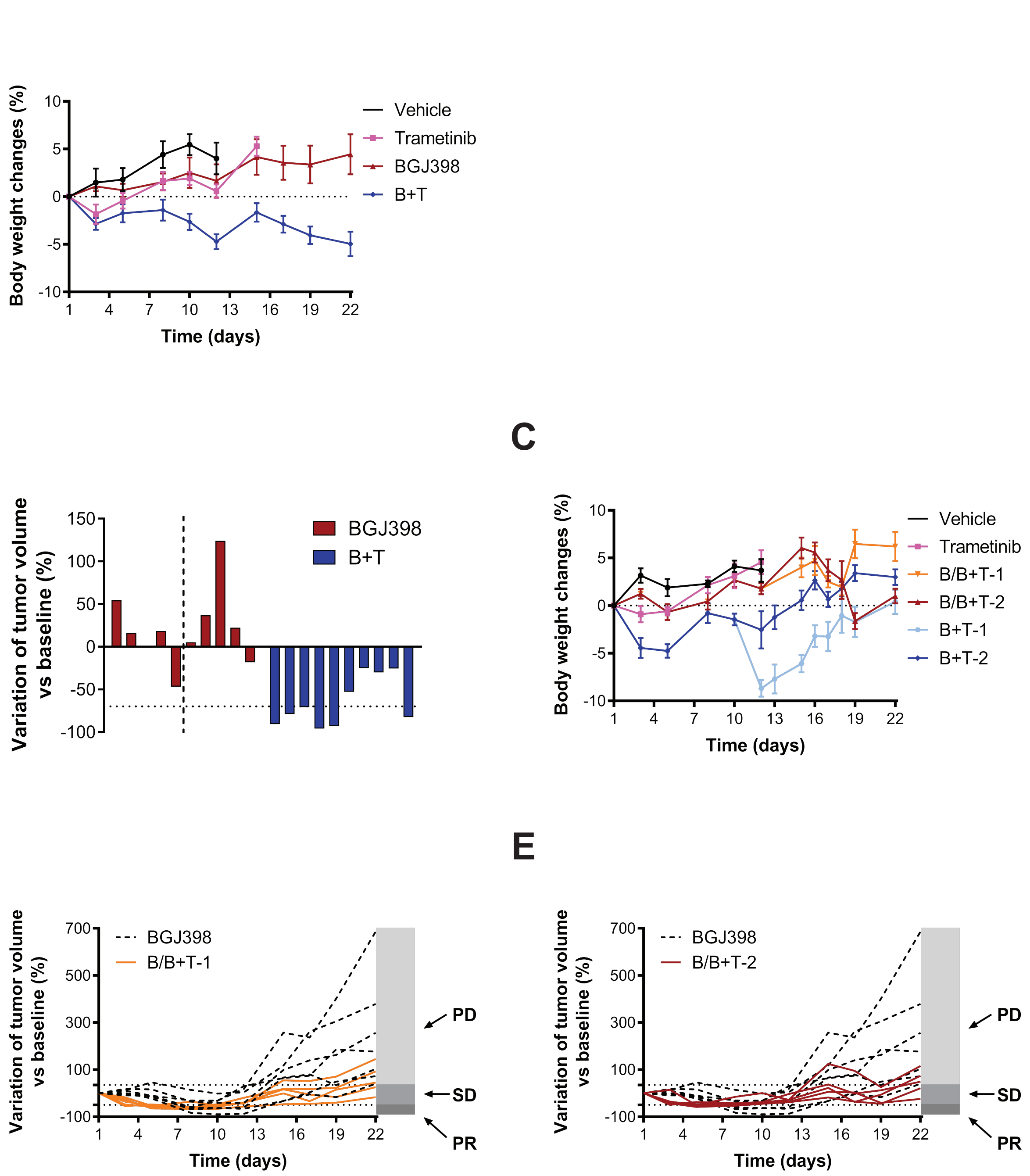

### Supplementary Figure 11

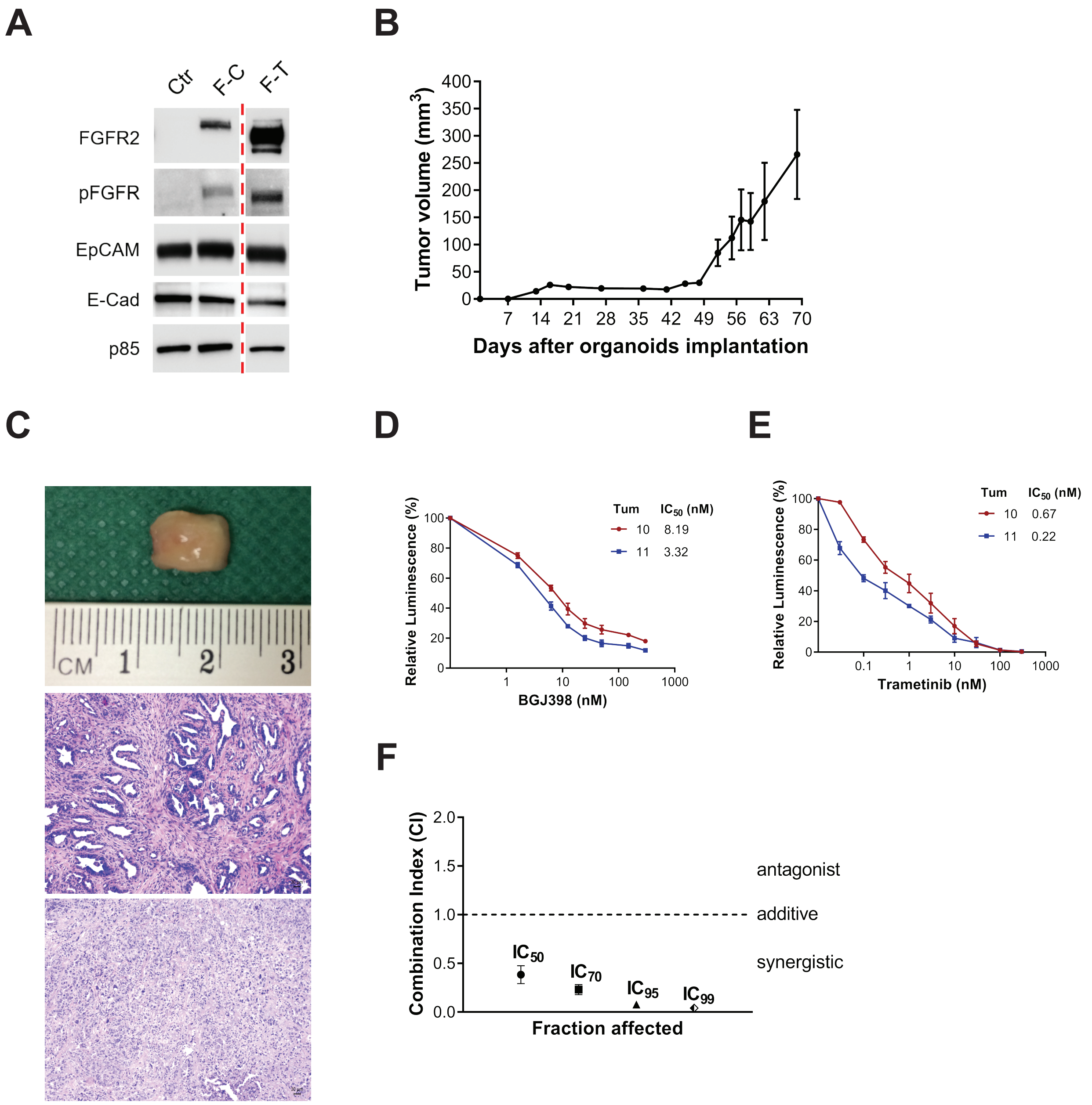

| GROUPS | INTRAHEPATIC | SUBCUTANEOUS |
|---|---|---|
| **Ctr** | 0/6 | 0/8 |
| **F-BICC1** | 4/4 | 4/4 |
| **F-MGEA5** | 0/4 | 1/8 |
| **F-TACC3** | 7/7 | 4/4 |
| **F-TACC3 V565F** | 2/2 | 4/4 |
| **F-CCDC6** | n.d. | 6/8 |

**Supplementary Table 1.** Tumorigenicity of genetically modified organoids upon transplantation in NOD-SCID mice. n.d., not done.

| Up-regulated in FF+ vs FF- | | |
|---|---|---|
| **NAME** | **NES** | **FDR** |
| INTERFERON ALPHA RESPONSE | 2,09 | 0 |
| TGF BETA SIGNALING | 2,03 | 0,001 |
| EPITHELIAL MESENCHYMAL TRANSITION | 1,94 | 0,002 |
| TNFA SIGNALING VIA NFKB | 1,84 | 0,002 |
| HYPOXIA | 1,70 | 0,007 |
| INTERFERON GAMMA RESPONSE | 1,67 | 0,008 |
| KRAS SIGNALING UP | 1,53 | 0,027 |

| Down-regulated in FF+ vs FF- | | |
|---|---|---|
| **NAME** | **NES** | **FDR** |
| XENOBIOTIC METABOLISM | -2,48 | 0 |
| FATTY ACID METABOLISM | -1,86 | 0,003 |
| BILE ACID METABOLISM | -1,85 | 0,002 |
| REACTIVE OXYGEN SPECIES PATHWAY | -1,62 | 0,016 |
| ADIPOGENESIS | -1,53 | 0,037 |
| PEROXISOME | -1,51 | 0,038 |

**Supplementary Table 2.** Gene set enrichment analysis (GSEA) results of FF+ and FF- iCCA samples from the TCGA cholangiocarcinoma dataset. Gene sets derive from MsigDB Hallmark collection. A positive Normalized Enrichment Score (NES) indicates gene sets up-regulated in FF+ patients whereas a negative NES indicates gene sets up-regulated in FF- patients. Only gene sets with False Discovery Rate (FDR) <0.05 are shown.

| REAGENTS AND CHEMICALS | SUPPLIER | IDENTIFIER |
|---|---|---|
| (Hydroxypropyl)methyl cellulose | Sigma-Aldrich | Cat #H7509 |
| [Leu15]-gastrin I human | Sigma-Aldrich | Cat #G9145 |
| Advanced DMEM/F-12 | Thermo Fisher Scientific | Cat #12634-010 |
| Agarose LE | Euroclone | Cat #EMR920500 |
| Albumin (BSA) Fraction V (pH 7.0) | PanReac AppliChem | Cat #A1391 |
| B27 supplement 50X minus vitamin A | Thermo Fisher Scientific | Cat #12587-010 |
| Collagenase from Clostridium histolyticum | Sigma-Aldrich | Cat #C9407 |
| Crystal Violet | Sigma-Aldrich | Cat #C0775 |
| Dimethyl Sulfoxide, Cell Culture Reagent | MP Biomedicals | Cat #02196055-CF |
| Dispase II | Thermo Fisher Scientific | Cat #17105-041 |
| DMEM F-12 | Lonza | Cat #12-719F |
| DMEM with Glucose and L-Glutamine | Lonza | Cat #12-604F |
| DMEM, high glucose, GlutaMAX, pyruvate | Thermo Fisher Scientific | Cat #31966-021 |
| DNaseI | Sigma-Aldrich | Cat #DN25 |
| EuroRT M-MLV Reverse Transcriptase | Euroclone | Cat #EMR438050 |
| Random Hexamers | Euroclone | Cat #EMR428200 |
| Foetal Bovine Serum (FBS) | Euroclone | Cat #ECS0180L |
| Gelatin from porcine skin | Sigma-Aldrich | Cat #G8150 |
| GeneRuler 100 bp Plus DNA Ladder | Thermo Fisher Scientific | Cat #SM0321 |
| Glutamax | Thermo Fisher Scientific | Cat #35050-068 |
| HEPES 1 M | Euroclone | Cat #ECM0180D |
| Infigratinib (BGJ398) | MedChemExpress | Cat #HY-13311 |
| L-Glutamine | Lonza | Cat #17-605E |
| Lipofectamine 3000 Transfection Reagent | Thermo Fisher Scientific | Cat #L3000015 |
| Matrigel matrix, phenol-red-free | Corning | Cat #356231 |
| Matrigel matrix | Corning | Cat #354234 |
| N2 supplement 100X | Thermo Fisher Scientific | Cat #17502-048 |
| N-acetyl-L-cysteine | Sigma-Aldrich | Cat #A9165 |
| Nicotinamide | Sigma-Aldrich | Cat #N0636 |
| NP-40 Surfact-Amp Detergent Solution | Thermo Fisher Scientific | Cat #28324 |
| Paraformaldehyde (PFA) | Sigma-Aldrich | Cat #158127 |
| PBS-1X | Thermo Fisher Scientific | Cat #14190-094 |
| Penicillin/Streptomycin mixture | Lonza | Cat #17-602E |
| Poly(ethylene glycol) with 400 Average Mn | Sigma-Aldrich | Cat #202398 |
| Polypropylen glycol P400 | Sigma-Aldrich | Cat #81350 |
| Prestained Protein SHARPMASS VII | Euroclone | Cat # EPS026500 |
| ProLong Gold Antifade Reagent with DAPI | Cell Signaling Technology | Cat #8961 |
| Proteinase K, recombinant PCR Grade | Roche | Cat #03115828001 |
| QIAzol Lysis Reagent | QIAGEN | Cat #1023537 |
| Recombinant human FGF10 | PeproTech | Cat #100-26 |
| Recombinant human HGF | PeproTech | Cat #100-39 |
| Recombinant human Noggin | PeproTech | Cat #120-10C |
| Recombinant murine EGF | PeproTech | Cat #315-09 |
| Recovery Cell Culture Freezing Medium | Thermo Fisher Scientific | Cat #12648-010 |

| | | |
|---|---|---|
| Rho kinase inhibitor Y-27632 dihydrochloride | Sigma-Aldrich | Cat #Y0503 |
| RNAlater RNA Stabilization Reagent | QIAGEN | Cat #76104 |
| Rspo1-conditioned medium | homemade | |
| SHP099 | MedChemExpress | Cat #HY-100388 |
| SYBR Green PCR Master Mix | Thermo Fisher Scientific | Cat #4309155 |
| Trametinib | MedChemExpress | Cat #HY-10999 |
| TRIzol Reagent | Thermo Fisher Scientific | Cat #15596026 |
| Triton X-100 Molecular Biology grade | PanReac AppliChem | Cat #A4975 |
| TrypLE Express | Thermo Fisher Scientific | Cat #12605-028 |
| Tween-20 | Fisher BioReagents | Cat #BP337-100 |
| Tween-80 | Sigma-Aldrich | Cat #P1754 |
| Wnt3a-conditioned medium | homemade | |
| Wonder Taq Hot Start | Euroclone | Cat #EME023500 |
| XenoLight D-Luciferin Potassium Salt | PerkinElmer | Cat #122799 |

**Supplementary Table 3.** List of reagents and chemicals.

| ANTIBODIES | SUPPLIER | IDENTIFIER | WORKING CONDITIONS | APPLICATION |
|---|---|---|---|---|
| Recombinant Anti-SHP2 (phospho Y542) [EP508(2)Y] | Abcam | Cat #ab62322 | 1:1000 | WB |
| FRS2 | Abcam | Cat #ab200548 | 0.2 µg/ml | WB |
| Recombinant Anti-Cytokeratin 19 [EP1580Y] | Abcam | Cat #ab52625 | 1:20000 | WB |
| | | | 1:400 | IF |
| | | | 1:400 | IHC |
| Recombinant Anti-EpCAM [EPR20533-63] | Abcam | Cat #ab221552 | 1:5000 | WB |
| E-cadherin | BD bioscience | Cat #610181 | 1:5000 | WB |
| Goat Anti-Mouse IgG (H + L)-HRP Conjugate | Bio-Rad | Cat #170-6516 | 1:10000 | WB |
| Goat Anti-Rabbit IgG (H + L)-HRP Conjugate | Bio-Rad | Cat #170-6515 | 1:20000 | WB |
| Phospho-FGF Receptor (Tyr653/654) (55H2) | Cell Signaling Technology | Cat #3476 | 1:1000 | WB |
| SHP2 | Cell Signaling Technology | Cat #3752 | 1:1000 | WB |
| Phospho-FRS2-α (Tyr196) | Cell Signaling Technology | Cat #3864 | 1:1000 | WB |
| p44/42 MAPK (Erk1/2) | Cell Signaling Technology | Cat #9102 | 1:1000 | WB |
| Phospho-p44/42 MAPK (Erk1/2) (Thr202/Tyr204) (D13.14.4E) | Cell Signaling Technology | Cat #4370 | 1:1000 | WB |
| | | | 1:200 | IHC |
| MEK1/2 (L38C12) | Cell Signaling Technology | Cat #4694 | 1:1000 | WB |
| Phospho-MEK1/2 (Ser217/221) (41G9) | Cell Signaling Technology | Cat #9154 | 1:1000 | WB |
| PI3 Kinase p85 | Cell Signaling Technology | Cat #4292 | 1:1000 | WB |
| Cleaved PARP (Asp214) (D6X6X) | Cell Signaling Technology | Cat #94885 | 1:1000 | WB |
| Cyclin D1 (E3P5S) | Cell Signaling Technology | Cat #55506 | 1:1000 | WB |
| Ki-67 (D3B5) | Cell Signaling Technology | Cat #12202 | 1:200 | IHC |
| Hepatocyte (Concentrate) HepPar1 | Dako | Cat #M7158 | 1:100 | IHC |
| Fluorescein (FITC) AffiniPure Goat Anti-Mouse IgG (H+L) | Jackson ImmunoResearch | Cat #115-095-003 | 1:100 | IF |
| Rhodamine (TRITC) AffiniPure Goat Anti-Rabbit IgG (H+L) | Jackson ImmunoResearch | Cat #111-025-045 | 1:100 | IF |
| Albumin | Novus Biologicals | Cat #NB600-41532 | 1:20000 | WB |

| | | | | |
|---|---|---|---|---|
| MYC 9E10 mouse hybridoma | Purified in house | | 2 µg/ml | WB |
| | | | 2 µg/ml | IF |
| | | | 1.6-3.2 µg/ml | IHC |
| β-Actin | Sigma-Aldrich | Cat #A1978 | 1:20000 | WB |

**Supplementary Table 4.** List of antibodies.

| PRIMERS | Sequence (5'--------3') | SIZE amplicon | Tm |
|---|---|---|---|
| **mHprt_F** | AAG CTT GCT GGT GAA AAG GA | 186 bp | 55.2°C |
| **mHprt_R** | TTG CGC TCA TCT TAG GCT TT | | |
| **mLgr5_F** | TGA CTT TGA ATG GTG CCT CG | 176 bp | 60.3°C |
| **mLgr5_R** | GGG TAA GTC TTC GAG TAG GTT G | | |
| **mKrt7_F** | ACG TCA AAG CCC AGT ATG AG | 147 bp | 57.9°C |
| **mKrt7_R** | TCA TCT CCG CAA TCT CAT TCC | | |
| **mKrt19_F** | CCA GGA AGC CCA CTA CAA CAA T | 117 bp | 60.3°C |
| **mKrt19_R** | TCG AGG GAG GGG TTA GAG TAA A | | |
| **mTtr_F** | ATG GTC AAA GTC CTG GAT GC | 233 bp | 57.9°C |
| **mTtr_R** | AAT TCA TGG AAC GGG GAA AT | | |
| **mHnf4α7_F** | GGG TAC CCT TGG TCA TGG TCA GTG | 334 bp | 66.1°C |
| **mHnf4α7_R** | GCT TCC TTC TTC ATG CCA GCC CGG | | |
| **mG6p_F** | GAA TTA CCA AGA CTC CCA GG | 572 bp | 57.3°C |
| **mG6p_R** | TGA GAC AAT ACT TCC GGA GG | | |
| **mCyp3a_F** | TGG TCA AAC GCC TCT CCT TGC TG | 105 bp | 64.2°C |
| **mCyp3a_R** | ACT GGG CCA AAA TCC CGC CG | | |
| **mCftr_F** | CAC AGA CCT CAT TGC CTC AC | 104 bp | 59.4°C |
| **mCftr_R** | CCT CAA AAT TGG TGT GGT CC | | |
| **mActb_F** | TGA CAG GAT GCA GAA GGA GA | 82 bp | 57.3°C |
| **mActb_R** | GTA CTT GCG CTC AGG AGG AG | | |

**Supplementary Table 5.** List of primers.

| KITS | SUPPLIER | IDENTIFIER |
|---|---|---|
| CellTiter-Glo Luminescent Cell Viability Assay | Promega | Cat #G7571 |
| Masson's trichrome kit | Diapath | Cat #010210 |
| miRNeasy Mini Kit | Qiagen | Cat #217004 |
| PEG Virus Precipitation Kit | Abcam | Cat #ab102538 |
| Pierce BCA Protein Assay kit | Thermo Fisher Scientific | Cat #23227 |
| TruSeq Stranded mRNA Kit | Illumina | Cat #20020594 |

**Supplementary Table 6.** List of kits.
